# PartiNet is a dynamic adaptive neural network for high-performance particle picking in cryo-electron microscopy

**DOI:** 10.64898/2026.01.23.700950

**Authors:** Mihin Perera, Winnie Tan, Edward Yang, Ojasvi Jain, Mansi Aggarwal, Hariprasad Venugopal, Julie Iskander, Joseph D. Berry, Andrew Leis, Shabih Shakeel

## Abstract

Accurate, efficient and autonomous particle picking is a major bottleneck in high-resolution cryo-electron microscopy (cryo-EM). We introduce PartiNet, an Artificial Intelligence (AI)-based particle picker with size-agnostic detection and pre-trained models that eliminate manual parameter specification and dataset-specific training, pioneering dynamic neural network inference for single particle cryo-EM pipeline. Unlike static architectures, PartiNet employs a dynamic framework that adjusts network complexity in real-time based on perceived micrograph quality. This adaptive approach accelerates inference up to 7-fold compared to existing tools without sacrificing particle selection quality. Training on diverse protein datasets showed that PartiNet improves particle yields, enhances sampling of rare orientations, and is compatible with on-the-fly workflows. Comprehensive evaluation on benchmark datasets and validation on a new dataset of the chromatin remodeler MORC2 demonstrates superior precision and recall, with the ability to detect heterogeneous protein species, leading to more complete structural models and consistently higher-resolution reconstructions.

## Introduction

Three-dimensional (3D) reconstructions of protein molecules from cryo-EM data enable the generation of atomically detailed chemical models that underpin studies of normal and pathological biological function and guide therapeutic development^1^. These reconstructions are obtained by advanced image processing methods that combine large numbers of two-dimensional (2D) images containing individual protein “particles” extracted from electron micrographs, often achieving resolutions better than ∼4 Å, where secondary structure becomes unambiguous^1,2^. Cryo-EM micrographs are inherently noisy, low-dose projections of ensembles of protein molecules, making accurate identification of individual particles a critical early step^2^. During particle picking, candidate particles are detected, extracted, and subsequently aligned and averaged to generate a 3D reconstruction^3,4^. This step is particularly challenging due to low signal-to-noise ratios, conformational heterogeneity, contaminating features, and the scale of modern cryo-EM datasets. Consequently, particle detection remains a major bottleneck affecting both the accuracy and throughput of cryo-EM structure determination^5,6^.

Classical particle picking relied on heuristic approaches like template matching and feature-based detection. In template matching, users manually select ‘representative’ particles as templates, which are searched across micrographs dataset using the Cross Correlation Function (CCF)^7,8^, with correlation peaks indicating candidate particles. Feature-based methods instead apply image filters to detect regions of rapid intensity change, most commonly using the Laplacian of Gaussian (LoG) filter^9–11^. However, LoG performance is sensitive to filter parameters and often struggles with low contrast or overlapping particles. Both CCF- and LoG-based methods are also prone to false positives from ice contaminants (frost), which can produce intense signal peaks due to high electron beam opacity^11^. These limitations prompted the development of AI-based methods for particle detection.

Particle detection can be framed as an AI object-detection problem, in which each particle coordinate (object localisation) and boundary of each particle (object segmentation) must be determined accurately to extract the particle for subsequent 3D reconstruction. A number of AI-based particle picking programs have been developed, including DeepPicker^12^, APPLE^13^, Topaz^14^, crYOLO^15^, PIXER^16^, CASSPER^17^, CryoTransformer^18^ and CryoSegNet^6^, with Topaz and crYOLO among the most widely used. Topaz is a Convolution Neural Network (CNN)-based picker that frames the task as a positive-unlabeled (PU) learning problem^14^. This approach requires prior identification of a small number of positive particle regions^19,20^. crYOLO adapts the YOLO9000 object detection framework^21^; it uses image tiling to divide images into grids, and predicts the presence of particle centres within each grid^15^.

Whilst AI methods have improved particle picking accuracy and throughput, important limitations remain. Firstly, current methods struggle to detect particles in challenging micrographs with complex noise profiles, contamination, ice-thickness gradients or sample heterogeneity^5,6^. Secondly, all current methods employ developer-defined static architectures, applying the same network capacity to all images regardless of complexity^22,23^. This one-size-fits-all approach is computationally inefficient for straightforward images and may be inadequate for challenging ones. Finally, with the exception of CryoTransformer and CryoSegNet, many models have been trained on relatively small datasets (<1000 micrographs), which can limit generalisability and often necessitates de novo training or fine-tuning of models for new data.

Two recent developments in machine learning and cryo-EM provide opportunities to overcome these limitations. The first is the advancement of dynamic architecture frameworks in machine learning, which allow model architectures to adapt during inference and detection, promising improved adaptability to varying image complexity^22–24^. To date, no dynamic AI particle picking algorithms have been developed for cryo-EM^25^. The second is the creation of CryoPPP^26^ - a comprehensive, curated, dataset of particle coordinates from EMPIAR specifically for machine learning applications. CryoPPP has already accelerated the development of new architectures, with CryoSegNet and CryoTransformer matching or outperforming Topaz and crYOLO in terms of final map resolution^6,18^.

Here, we introduce PartiNet, a novel particle picking method based on the DynamicDet neural network architecture^24^. We trained this framework on the CryoPPP dataset with custom denoiser pre-processing stages and demonstrate the efficacy of the dynamic architectures for protein structure determination. Through comprehensive benchmarking on test datasets and validation on full-scale experimental datasets including the chromatin remodeler MORC2, we show that PartiNet matches or exceeds the quality of particles extracted by current AI pickers while processing micrographs up to 7 times faster.

## Results

### PartiNet is a dynamic particle picker

PartiNet employs a dynamic architecture that adapts network complexity in response to micrograph difficulty, distinguishing it from existing, static particle-picking methods^24^ (Fig 1a). The system comprises two detectors and an adaptive router that analyses extracted features to classify micrographs relatively as ‘easy’ or ‘hard’. Based on this assessment, images are processed either by a single detector for straightforward cases or two cascaded detectors for perceived challenging micrographs, dynamically scaling computational resources to match image requirements *in situ* (Fig 1b). Our implementation utilises dual YOLOv7 detectors with a shallow convolutional neural network adaptive router^20,24^. PartiNet differs from the existing YOLO network for particle picking by crYOLO in two key respects: it leverages the newer YOLOv7 architecture rather than a customised YOLO9000 implementation, and it dynamically adjusts network depth based on the input micrograph instead of relying on a single static model. Together, these design choices enable PartiNet to flexibly balance accuracy and computational efficiency across heterogeneous cryo-EM datasets.

**Figure 1.**
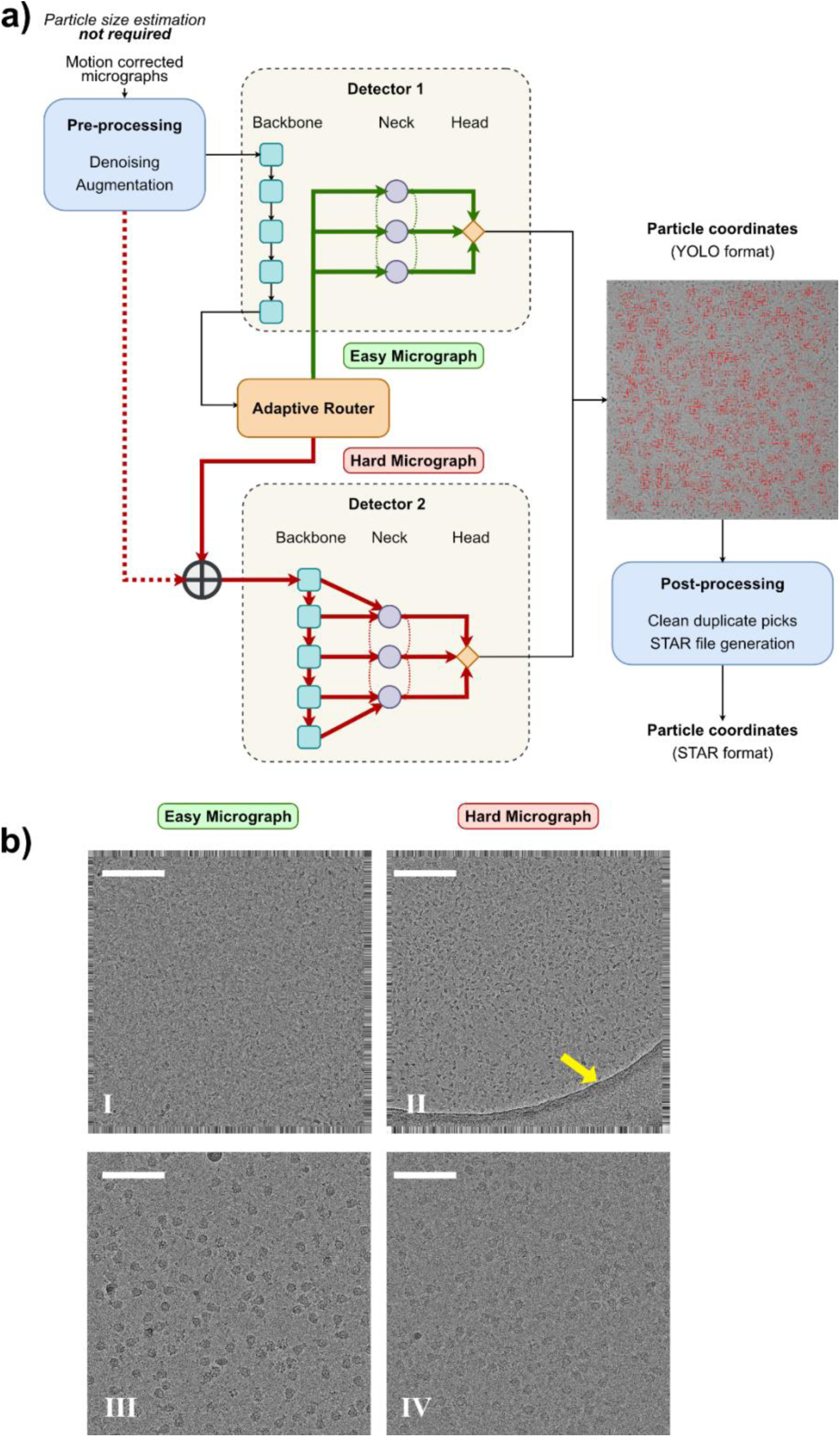
PartiNet has a dynamic architecture. **a.** PartiNet employs a dynamic architecture with dual detectors and an adaptive router. Micrographs are denoised and processed through Detector 1’s backbone to generate feature maps. The adaptive router assigns difficulty scores to these feature maps, directing easy micrographs back through Detector 1’s neck and head, while difficult micrographs are concatenated with the original image and processed through Detector 2. A post-processing module removes duplicate picks and converts YOLO coordinates to STAR format. **b.** The adaptive router differentiates micrographs based on imaging conditions without supervision. (I-II) Two micrographs from EMPIAR-10017 showing PartiNet’s classification of the micrograph containing support film (yellow arrow) as more difficult than the one without support film. Fringing artifacts from aggressive motion correction are visible at image edges. Scale bar, 100 nm. (III-IV) Two micrographs from EMPIAR-10089 demonstrating PartiNet’s identification of the lower defocus micrograph as more difficult due to reduced (phase) contrast. The difficult micrograph is at -1.07 µm defocus versus -2.07 µm for the easy micrograph. Scale bar, 120 nm.

### PartiNet enables turnkey particle picking without size specification or training

PartiNet enables truly out-of-box particle picking through a simple, single-command interface that requires no parameter specification, model training, or manual intervention. Unlike existing methods that require users to specify expected particle size, train on their own datasets, or manually adjust detection parameters, PartiNet automatically detects particles across all sizes using pre-trained models. The software accepts raw motion-corrected micrographs as input and outputs particle coordinates with confidence scores, integrating seamlessly into automated cryo-EM workflows. This is achieved through a modified Python^27^ implementation of DynamicDet^24^, built with PyTorch^28^, expanded to read motion-corrected MRC micrographs and output particle coordinates in STAR file format compatible with popular processing packages (Fig 1). We have provided a custom Wiener filter-based denoiser algorithm, based on the work of CryoSegNet and CryoTransformer^6,18^, with a novel multiprocessing layer allowing for efficient denoising of large datasets. We have also provided scripts for preparing custom datasets for training PartiNet on user-generated datasets and for finetuning of the current model. PartiNet is designed for command line use and supports high performance computing (HPC) environments, enabling scalable multi-GPU/CPU execution via job schedulers such as Slurm. Typical PartiNet workflows comprises three automatable steps: micrograph denoising, particle detection, and coordinate filtering with conversion to STAR format for downstream analysis (Fig 1). The ability to perform size-agnostic particle picking, unlike crYOLO and Topaz, represents a key feature of PartiNet and enables seamless integration into fully automated, unsupervised cryo-EM image processing pipelines.

### PartiNet training on comprehensive dataset enables generalisation without user-specific fine-tuning

PartiNet was trained using the CryoPPP dataset, a curated benchmark comprising ∼10,000 labelled micrographs from 34 cryo-EM protein datasets (Supplementary Table 1). These datasets span diverse protein sizes, symmetries and imaging conditions, and include gold standard particle coordinates manually curated by the authors of CryoPPP^29^, enabling PartiNet to generalise to new datasets without requiring user-specific training or fine-tuning - a significant advantage over methods that require retraining for optimal performance. We used seven, randomly selected datasets as test sets for the comparison of crYOLO, Topaz and PartiNet performance. For the remaining datasets, 80% of micrographs from each dataset were allocated randomly for training, and the remaining 20% were used for validation. The training, validation and test sets comprised 6224, 1563 and 1879 micrographs, respectively. Since each of the CryoPPP datasets selected contained only 300 micrographs per set, we also prepared 5 full size datasets for particle picking and reconstruction: MORC2 bound with the H3K9me3 peptide (in-house), rabbit muscle aldolase previously tested with Topaz (EMPIAR-10215)^14^, TcdA1 originally published with crYOLO (EMPIAR-10089)^15^, and two heterogeneous samples: Ankyrin-1 complex (EMPIAR-11043)^30^ and MlaCD complex (EMPIAR-12531)^31^. This design enabled systematic assessment of performance across diverse sample sizes and levels of heterogeneity.

### PartiNet preprocesses micrographs for enhanced picking

An obvious first step in particle picking from noisy cryo-EM data is some form of denoising. During PartiNet testing, we observed qualitative improvements for precision and recall on denoised micrograph datasets. To investigate this, we compared PartiNet’s performance across four preprocessing conditions: raw motion-corrected micrographs, and motion-corrected micrographs processed with three different denoising methods: two popular deep learning-based denoisers Janni^32^ and Topaz^33^ (both using theNoise2Noise framework^34^), and a heuristic Wiener filter-based algorithm introduced in CryoSegNet^6^.

Training was conducted for 100 epochs using CryoPPP datasets across all four conditions. We evaluated the pre-trained models provided by the Topaz and JANNI developers to assess their out-of-the-box performance, rather than training custom denoiser models for each optical condition in our dataset. We acknowledge that users may see improved performance with Topaz and JANNI denoisers using trained sub-models^33^.

For each training condition, we calculated and plotted the Mean Average Precision @ 50% confidence, Precision, and Recall (Supplementary Fig 1a). We observed that PartiNet-CryoSegNet denoise scored the highest across these metrics, followed closely by raw motion-corrected micrographs and then Topaz-denoised, with JANNI-denoised data lagging substantially across all metrics.

We suspected that JANNI and Topaz had sub-par performance in our testing due to the shared underlying Noise2Noise algorithm adversely affecting the quality of the particles in the micrographs. We plotted the particle coordinates generated by PartiNet for all 4 regimes on the same micrograph selected from the Influenza haemagglutinin trimer dataset (EMPIAR-10093; Supplementary Fig 1b-d). With no denoising applied, PartiNet was able to identify some particles but struggled with the complex noise of the image, as well as differentiating individual particles near contamination (Supplementary Fig 1b). We observed that both Topaz and JANNI suppressed high frequency information across this micrograph and seemed to be especially susceptible to this behaviour when electron-dense contamination was present in the micrograph. Particles close to contamination were “flattened”, preventing effective delineation of closely packed particles, both during denoising and subsequent picking with PartiNet (Supplementary Fig 1c-d). Conversely, the PartiNet implementation of CryoSegNet denoise retained discrete particles present in the micrographs, even when the particles were closely packed to the boundary of signal-dense contamination (Supplementary Fig 1e).

Given the efficacy of the CryoSegNet denoiser in conjunction with PartiNet, we integrated it directly into our software. The original denoiser was a single-threaded process, significantly bottlenecking denoising of large datasets. We developed a multiprocessing pipeline for this denoiser using the *concurrent.futures* module in Python, allowing for asynchronous execution of batch denoising for micrographs. Our implementation is especially suited for efficient use of HPC resources using job schedulers. Denoising 1000 micrographs on a single node with 32 CPU cores had a CPU efficiency of 94.47% for 01:41:20 (hh:mm:ss) walltime, thus demonstrating that denoising can be performed efficiently at scale without introducing a preprocessing bottleneck.

During training, we augmented each micrograph before passing it through PartiNet. Augmentations comprised a random combination of image transformations, each of which had a specified probability to be applied to every micrograph during each epoch (Supplementary Table 2, Supplementary Fig 2). Across training epochs, micrographs were randomly augmented for each pass. This served to prevent model overfitting and increased the feature space of the training data for PartiNet to learn ^35^. During inference, PartiNet employs test-time augmentation (TTA) to improve detection robustness and accuracy ^20^. Each micrograph is processed at multiple scales (100%, 83%, 67%) and with horizontal flipping to increase the feature space available for picking. The predictions from all augmented versions are then transformed back to the original coordinate system and combined. This allows PartiNet to aggregate features identified at different resolutions and orientations, increasing detection confidence through multiple perspectives of the same micrograph.

### PartiNet implements a refined detector configuration

PartiNet was designed with YOLOv7 networks for each detector. We prepared 6 different configurations of YOLOv7^20^, with differing numbers of model parameters, input sizes, reported inference speed in image frames per second, and reported performance on the benchmarking object detection dataset COCO in mAP@50% (Supplementary Table 3)^36^. We trained each of these configurations for 100 epochs on CryoPPP training data denoised with PartiNet-CryoSegNet. We measured the mAP@50%, Precision and Recall, and observed that for all metrics, YOLOv7-W6 had the highest scores after 100 epochs of training (Supplementary Fig 3). YOLOv7, YOLOv7-X and YOLOv7-E6E achieved similar performance metrics, ranking as the second-best performing group, withYOLOv7-E6 scoring the lowest overall. In our hands, YOLOv7-D6 failed to converge, and would crash repeatedly during testing, and so was discontinued. Based on these results, we opted to use YOLOv7-W6 for all subsequent testing.

### Benchmarking PartiNet against established AI pickers shows improved performance

To compare the performance of PartiNet against Topaz, and crYOLO, we picked particles and reconstructed density maps on the 7 test datasets from cryoPPP (Supplementary Table 4). We evaluated the performance of PartiNet against gold-standard networks crYOLO and Topaz in terms of number of particles picked and the final resolution of the 3D maps. We mirrored a test workflow previously outlined in CryoSegNet and CryoTransformer to fairly compare the networks for all datasets^6,18^ (Supplementary Fig 4). We report the number of particles picked by each of Topaz, crYOLO and PartiNet, the number of particles remaining after “Select 2D” for reconstruction, and the final global resolution of the 3D map reconstruction with these selected particles (Fig 2).

**Figure 2.**
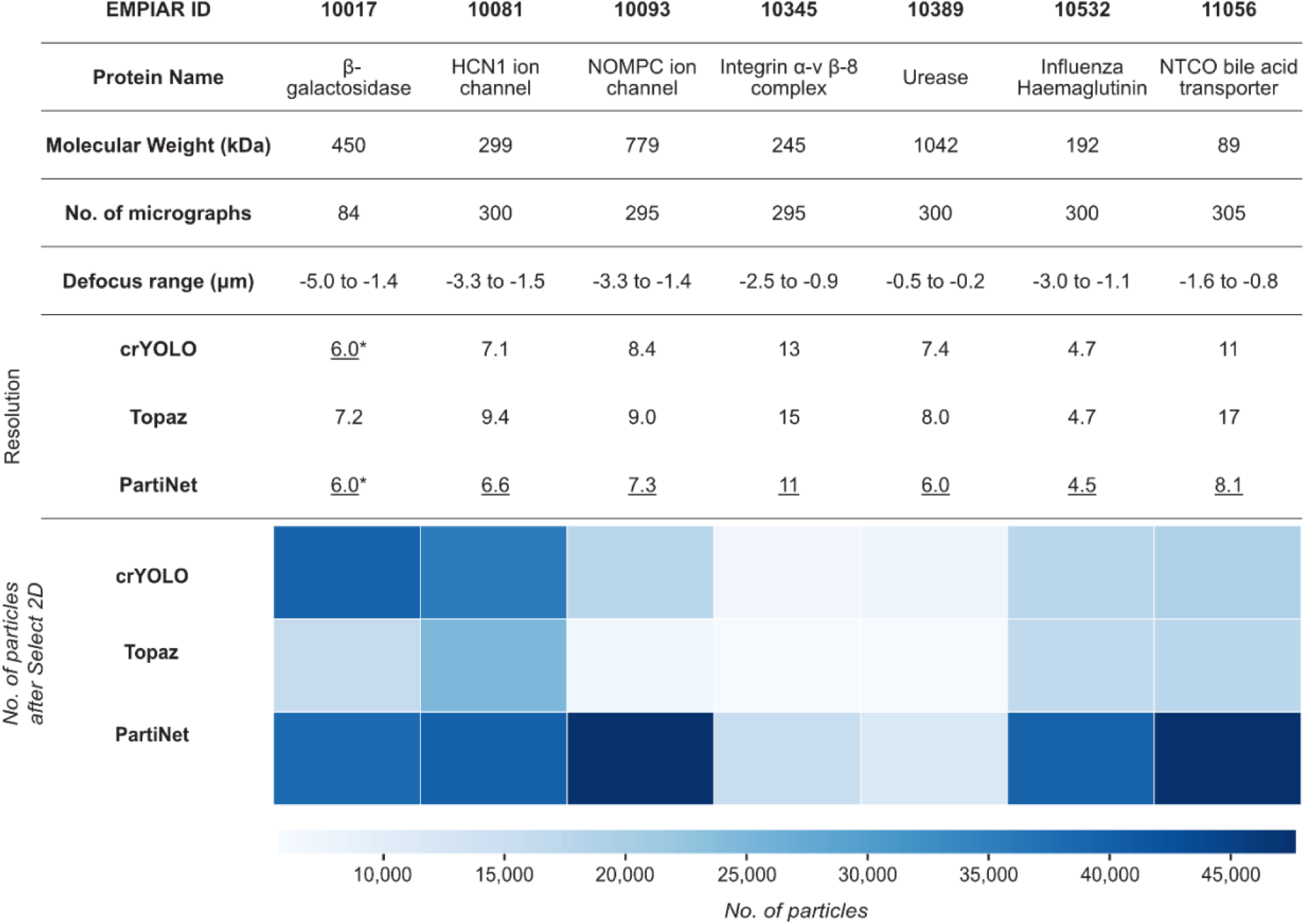
PartiNet outperforms popular particle pickers on small datasets. Seven datasets from CryoPPP were randomly selected to benchmark PartiNet against crYOLO and Topaz. The table summarises key dataset characteristics including EMPIAR ID, molecular weight, micrograph count, defocus range, and final map resolution, with the highest resolution result underlined for each protein. The accompanying heatmap shows the number of protein particles retained after 2D classification (“Select 2D”) for each method, with darker blue indicating higher particle numbers used for reconstruction.

For six of the seven test sets, PartiNet identified more particles prior to “Select 2D” filtering than the other AI methods, with crYOLO and Topaz alternating as the second-best performer. The exception was EMPIAR-10017, where crYOLO identified 3,000 more particles than PartiNet, with Topaz picking less particles (16,000) than both PartiNet and crYOLO. PartiNet excelled with smaller, lower molecular weight proteins (EMPIAR-10532 and EMPIAR-11056), where it substantially outperformed both crYOLO and Topaz in particle counts. This likely reflects PartiNet’s more aggressive identification strategy when dealing with smaller proteins that are inherently difficult to detect. A similar pattern was observed for EMPIAR-10093, despite this protein’s high molecular weight of 779 kDa. We attribute this to the challenging non-globular shape of NOMPC ion channel proteins, which complicates particle identification^14^. After “Select 2D” filtering, PartiNet consistently retained the most particles across all test sets except EMPIAR-10017, followed by crYOLO and then Topaz. Most importantly, PartiNet achieved the best final resolution for all datasets tested (Supplementary Fig 5). For EMPIAR-10017, this superior resolution was achieved despite PartiNet having fewer retained particles than crYOLO, suggesting that PartiNet identified higher-quality particles with better sampling of protein views. In fact, for all CryoPPP test sets, PartiNet demonstrated stronger sampling of protein views, with alignments of particles covering a broader range of Euler sphere when compared to crYOLO and Topaz (Supplementary Fig 6).

We extended our comparison of PartiNet, crYOLO and Topaz with two further datasets: TcdA1 (EMPIAR-10089)^15^ and rabbit muscle aldolase (EMPIAR-10215)^14^, published with crYOLO and Topaz method papers^14,15^, respectively. We first evaluated the computational efficiency of PartiNet and crYOLO by measuring picking speed in micrographs per second using per-micrograph inference timing (Fig 3a). Topaz was excluded from this speed comparison as it supports only single GPU processing, while both crYOLO and PartiNet support parallel processing with up to 4 GPUs under identical hardware conditions. PartiNet demonstrated substantially faster processing speeds than crYOLO across both datasets, with performance improvements of up to seven-fold on the rabbit muscle aldolase dataset (EMPIAR-10215) and a peak inference speed of 8 micrographs/second for TcdA1.

**Figure 3.**
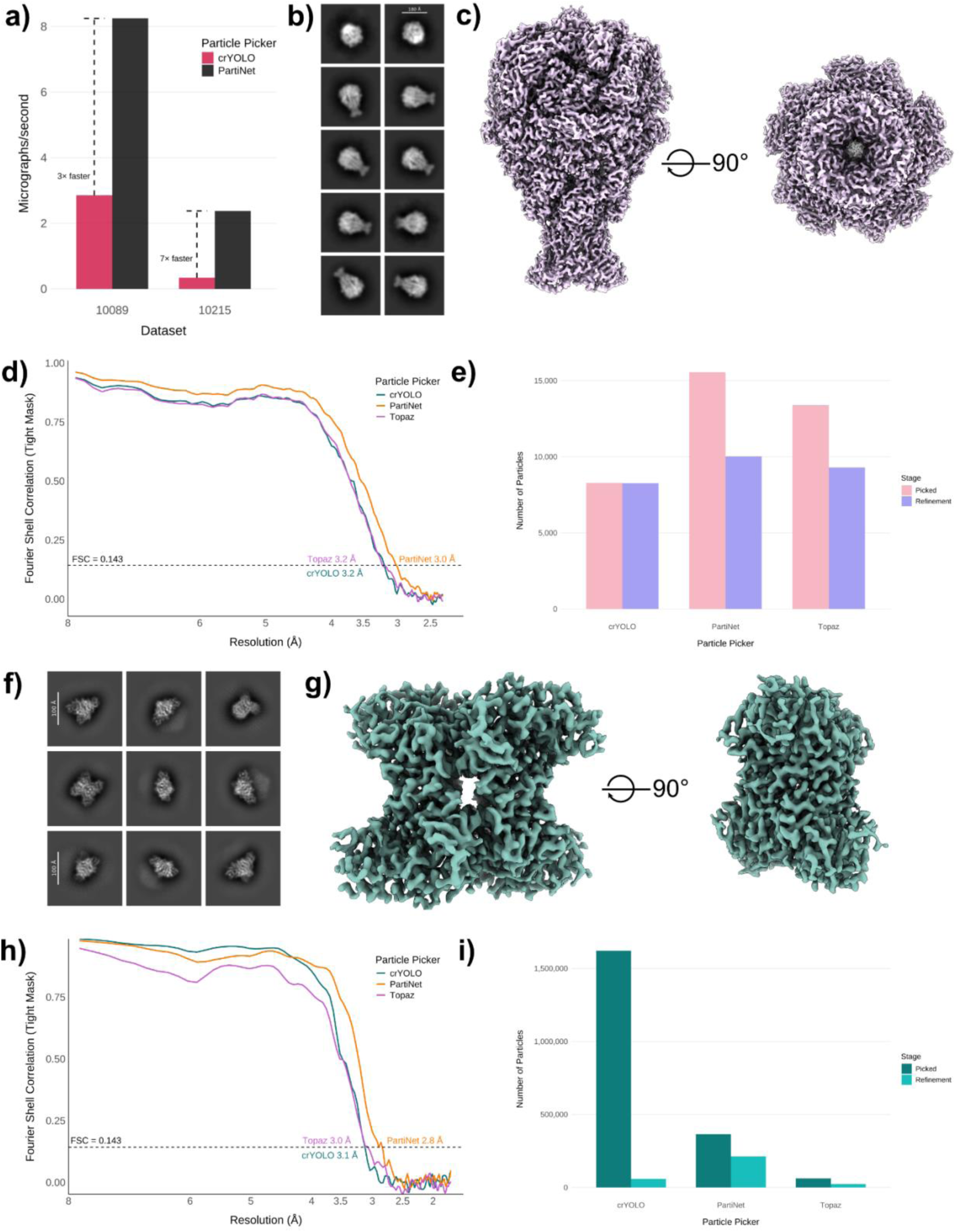
PartiNet outperforms other particle pickers on EMPIAR-10089 and EMPIAR-10215. **a.** Bar chart comparing picking speed of PartiNet (black) and crYOLO (red) on TcdA1 (EMPIAR-10089) and rabbit muscle aldolase (EMPIAR-10215), reported as micrographs per second. Approximate X-fold increase in speed with PartiNet is indicated for each dataset. **b.** Selected 2D class averages of TcdA1 particles picked with PartiNet. **c.** Cryo-EM map reconstruction of TcdA1 from PartiNet particles picked on EMPIAR-10089. **d.** FSC curves for final reconstructions of TcdA1 from particles picked with crYOLO (teal), Topaz (purple) and PartiNet (yellow) with resolution annotated at FSC = 0.143 cutoff. **e.** Bar chart comparing numbers of particles picked (pink) and used for final reconstruction (purple) of TcdA1 for PartiNet, crYOLO, and Topaz. **f.** Selected 2D class averages of rabbit muscle aldolase particles picked with PartiNet. **c.** Cryo-EM map reconstruction of rabbit muscle aldolase from PartiNet particles picked on EMPIAR-10215. **d.** FSC curves for final reconstructions of rabbit muscle aldolase from particles picked with crYOLO (teal), Topaz (purple) and PartiNet (yellow) with resolution annotated at FSC = 0.143 cutoff. **e.** Bar chart comparing numbers of particles picked (dark teal) and used for final reconstruction (light teal) of rabbit muscle aldolase for PartiNet, crYOLO, and Topaz.

Next, we applied a standardised workflow for processing TcdA1 to compare PartiNet, Topaz and crYOLO objectively (Supplementary Fig 7). We performed 2D classification on PartiNet-selected particles to assess particle quality through visual inspection of 2D class averages (Fig 3b). The resulting averages showed coherent, well-centred representations of TcdA1 spanning multiple orientations, with clearly resolved secondary structure features. Subsequent 3D reconstruction using PartiNet particles yielded a high-quality density map with well-defined secondary structure and no observable anisotropy that would result from particle orientation bias (Fig 3c). For comparative analysis, we reconstructed maps of TcdA1 with crYOLO and Topaz coordinates and plotted Fourier Shell Correlation (FSC) curves for each density map (Fig 3d). At an FSC cutoff of 0.143, PartiNet had the highest global resolution of 3.0 Å, followed by Topaz and crYOLO with 3.2 Å. These resolutions represent improvements over the 3.4 Å structure published originally for EMPIAR-10089, likely reflecting differences in reconstruction algorithms between SPHIRE and CryoSPARC. Finally, we compared PartiNet’s performance in terms of particle counts before and after cleaning by 2D classification (Fig 3e). PartiNet identified more particles (15,551) than Topaz or crYOLO (13,380 and 8,282, respectively). After filtering, PartiNet again retained the most particles (10,016) compared to crYOLO (9,292) and Topaz (8,258), revealing that PartiNet coordinates were well ranked. Interestingly, crYOLO retained effectively all particles picked, suggesting that the publicly available model for crYOLO may have been trained on this dataset prior to publication.

To complete analysis of TcdA1, we performed de novo, automated model building with the reconstructed maps from PartiNet, crYOLO and Topaz particles using ModelAngelo^37^ (Supplementary Fig 8). All three models demonstrated excellent sequence coverage and high ModelAngelo confidence scores for the majority of residues. Mean confidence scores were almost identical: PartiNet’s model had the highest mean confidence of 99.4%, followed by crYOLO with 99.2% and then Topaz with 98.9%. We also calculated the Root Mean Square Deviation (RMSD) in Å between a single monomer of the ModelAngelo prediction and the crystal structure of TcdA1 (PDB 4O9Y). PartiNet had the lowest RMSD of 0.67, followed by Topaz with 0.71 and then crYOLO with 0.78. All values were < 1.0 Å for most of the monomer sequence, indicating that resolutions of 3.0-3.2 Å are sufficient for accurate model building of TcdA1 in this dataset. We completed our assessment of the models by calculating Q-scores, an independent metric for map-model fitness, defined as the measure of an atom’s resolvability within a cryo-EM map. PartiNet had the highest Q-score with 0.68, followed by Topaz and crYOLO with 0.65, indicating PartiNet’s reconstruction led to the highest quality map built with ModelAngelo.

We repeated this analysis on rabbit muscle aldolase, again with a standardised workflow of picking with PartiNet, Topaz or crYOLO (Supplementary Fig 9). Subsequent 3D reconstruction of rabbit muscle aldolase with PartiNet showed a high-quality map with clear secondary structure, with no observable orientation bias or anisotropy (Fig 3g). At an FSC cutoff of 0.143, PartiNet had the highest global resolution at 2.8 Å, followed by Topaz with 3.0 Å and crYOLO with 3.1 Å (Fig 3h). We report the number of picked particles and those retained after filtering (Fig 3i). crYOLO picked the highest number of particles (1,621,138) compared to PartiNet (364,185) and Topaz (61,822); however, after filtering, PartiNet retained the most particles (212,176) followed by crYOLO (57,639) and Topaz (23,078). The large discrepancy between crYOLO’s picked and retained particles may be due to the size of the protein or the high density of particles in micrographs.

We again used ModelAngelo to validate the reconstructed maps of rabbit muscle aldolase (Supplementary Fig 10). Whilst PartiNet and crYOLO’s models had comparable sequence coverage, ModelAngelo was unable to build most of the sequence into the Topaz map. We inspected the map of rabbit muscle aldolase built with Topaz particles and observed strong anisotropy and poor reconstruction of secondary structure, suggesting that even though the map had a global resolution of 3.0 Å, it was not sufficient to build an appropriate model. We reported the confidence of residue predictions, as RMSD, between the ModelAngelo predictions and the crystal structure of rabbit muscle aldolase (PDB 6ALD), and map-model fit with Q-scores. PartiNet had the highest mean confidence of residue predictions at 99.7% followed by crYOLO with 99.1% and Topaz with 95.3%. PartiNet had the lowest mean RMSD of 0.45, followed by crYOLO with 0.53 and Topaz with 5.3. PartiNet had the highest mean Q-score of 0.68, followed by crYOLO with 0.66 and Topaz with 0.63. PartiNet consistently outperformed both crYOLO and Topaz across all metrics in terms of models built from particles picked.

Together, these results demonstrate that PartiNet consistently delivers superior particle selection, faster inference, and higher-quality reconstructions and atomic models across diverse datasets, establishing improved accuracy/efficiency trade-offs relative to existing AI-based particle pickers.

### Reconstructions from PartiNet particles enable comprehensive mapping of protein sequences

After confirming PartiNet’s performance on the 7 test datasets from CryoPPP, rabbit muscle aldolase and TcdA1, we extended our analysis to an unpublished dataset for the chromatin remodeler MORC2^38^ bound to the H3K9me3 peptide (Supplementary Fig 11) and compared performance against Topaz and crYOLO (Fig 4). First, we compared the picking speed of PartiNet against crYOLO for the dataset and measured PartiNet to be 5-fold faster (Fig 4a). Next, we again performed a standardised workflow for processing MORC2 with PartiNet, Topaz and crYOLO picks (Supplementary Fig 12). We plotted 2D averages for MORC2 particles picked with PartiNet, which showed convergence to high quality, coherent 2D classes, showing secondary structure and evidence of strong sampling across multiple views of MORC2 (Fig 4b). Next, we plotted the FSC curves for each map and observed that the 3D map reconstructed with PartiNet picks had a global resolution of 2.3 Å, compared to 2.5 Å and 2.7 Å with crYOLO and Topaz, respectively (Fig 4c). To assess the quality of the 3D map reconstruction from PartiNet, Topaz and crYOLO picks, we took the final reconstruction of each map and estimated the resolution of each voxel (rather than a global resolution) with the “Local Resolution Estimation” function in CryoSPARC (Fig 4d). As expected, the central globular structure of all MORC2 maps were resolved best due to this being comparatively rigid and electron dense. Conversely, the two double coil (CC) domain arms extending from the central body of the MORC2 were not resolved. This was expected, as these domains are highly flexible because they are responsible for the DNA compaction activity of MORC2, and the presence and activation of these domains are evidenced in other works where 3D variability analysis showed this movement^38^. The PartiNet map’s resolution was isotropic, whereas crYOLO’s and Topaz’s maps displayed local resolution loss on the fringes of the protein, and in the case of Topaz, within the central structure of MORC2 itself. Additionally, when viewed from the bottom, Topaz’s map exhibited anisotropy in the form of “smearing” between the flat front faces, indicating poor sampling of the top and bottom views of MORC2 during particle picking (Fig 4d, bottom panel). We extended our analysis to measure the quality of sampling across all views of MORC2. We calculated cFAR and SCF values for each map with the “Orientation Diagnostics” in CryoSPARC, with higher cFAR and SCF scores corresponding to stronger sampling across all views of the proteins^39,40^. This value is effectively an estimation of “orientation bias” in the map, where certain views of the protein are heavily sampled and rarer views of the protein are undersampled. This occurs due to the tendency of many proteins to partition to the air-water interface of the protein solution directly prior to vitrification, and represents a major bottleneck for data analysis. PartiNet had the highest cFAR score of 0.27, followed by crYOLO with 0.11 and the Topaz with 0.03. Interestingly, PartiNet had a SCF score of 0.77 compared to crYOLO’s 0.78, whereas Topaz has a substantially different SCF of 0.65. These discrepancies reveal interesting details about the particles picked by PartiNet and crYOLO. cFAR is calculated by measuring the correlations of half-maps in specific viewing angles in Fourier space^39^, meaning that both the alignment and quality of signal in that specific alignment is necessary for a good score. Conversely, SCF is calculated by quantifying how the alignments of particles cover the Euler sphere without quantifying signal quality from particles^40^. Given that PartiNet map had the highest global resolution and cFAR score, it can be concluded that PartiNet particles contributed meaningful particle signal information from alignments, with good coverage over the viewing angles of the protein. crYOLO, on the other hand, had comparable numbers of particles spanning the Euler sphere but may have contained more poor particles or particles that contributed poorly to the assigned alignment. LowTopaz scores for cFAR, SCF and number of particles indicate that these particles were of lesser quality and exhibited strong orientation bias, with poor contribution to signal in many alignments.

**Figure 4.**
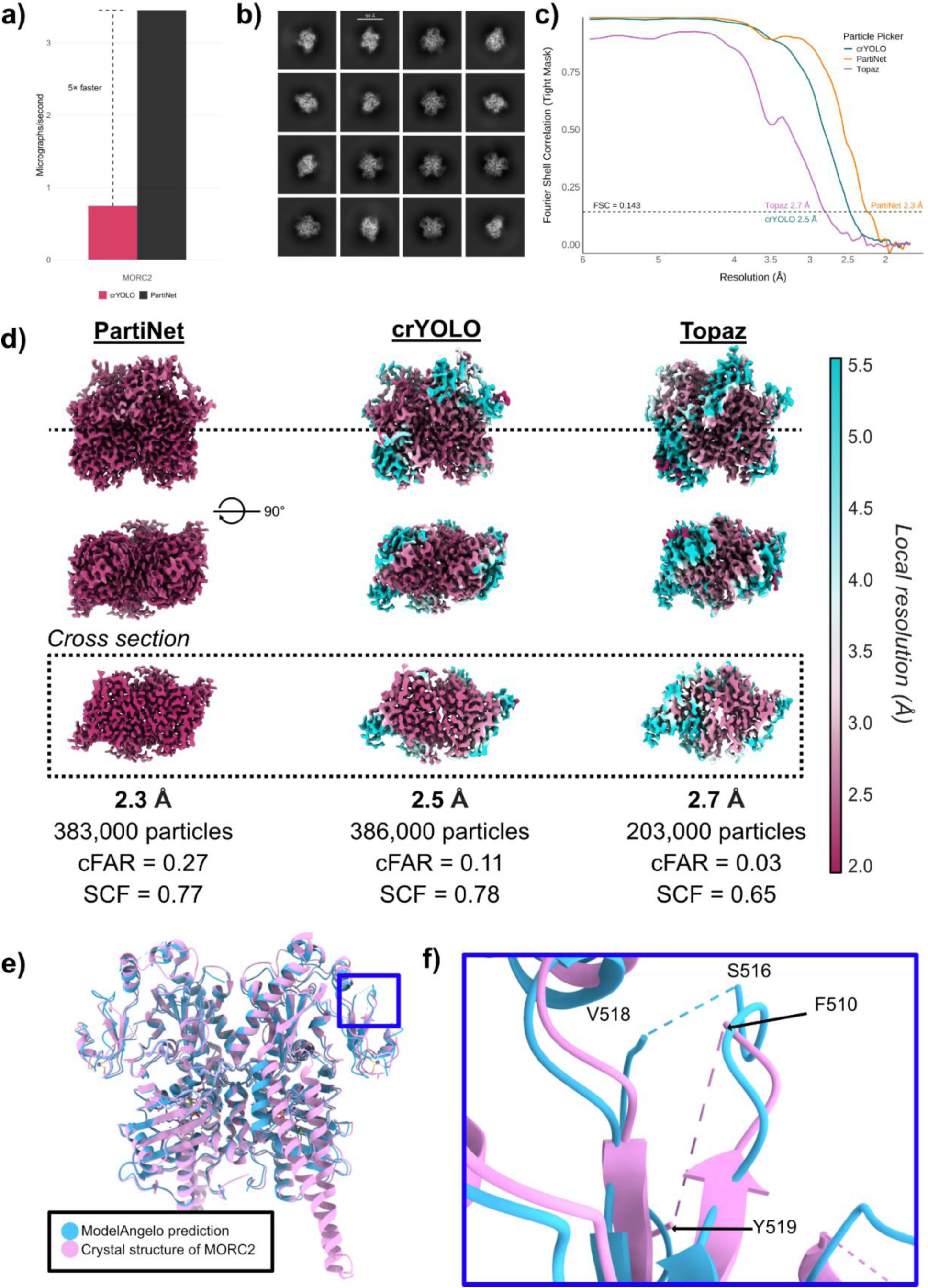
PartiNet picks allow high resolution reconstructions for model building. **a.** Bar chart comparing picking speed of PartiNet (black) and crYOLO (red) on MORC2. Approximate 5-fold increase in speed with PartiNet is indicated. **b.** Selected 2D class averages of MORC2 particles picked with PartiNet. **c.** FSC curves for final reconstructions of MORC2 from particles picked with crYOLO (teal), Topaz (purple) and PartiNet (yellow) with resolution annotated at FSC = 0.143 cutoff **d.** MORC2 maps were reconstructed using coordinates from PartiNet, Topaz, and crYOLO particle picking algorithms. Local resolution was estimated in CryoSPARC and visualised on the reconstructions in ChimeraX (v.1.10.1), where teal voxels indicate lower resolution and maroon indicates higher resolution. A central cross-section of each map is shown. The global resolution, final particle count, conical FSC area ratio (cFAR), and Sampling Compensation Factor (SCF) are shown for each map. **e.** The crystal structure of MORC2 (PDB 5OF9) was superimposed with the ModelAngelo-generated model from the PartiNet map after alignment in ChimeraX. The blue boxed region is shown in detail in **f.** highlighting differences in modeling of previously unresolved residues between the crystal structure and ModelAngelo prediction of MORC2.

We completed our analysis of MORC2 by using a reconstructed 3D map from PartiNet to build a structural model of MORC2 using ModelAngelo^37^. We superimposed this model with a published X-ray crystallography structure^41^ (PDB-5OF9, Fig 4e). We observed strong congruence between these models, especially in the central structure of the protein, with the notable absence of the two flexible coiled-coil (CC1) domains in our model. This is a limitation of cryo-EM, as highly flexible protein domains are difficult to resolve to high resolution with 3D cryo-EM maps without further advanced processing tools^42^. We plotted the per-residue confidence, RMSD with the crystal structure and Q-score for model-map fit (Supplementary Fig 13a-c). *In vitro* and *in vivo*, MORC2 exists as a homodimer^38,41^. We calculated RMSD for each monomer of each map, and found that our reconstructed map and the published map deviated within 1 Å of each other for the whole protein, except for unstructured tails and double CC domain arms, which are highly flexible (Supplementary Fig 13d).We observed a segment of the MORC2 protein that was mapped in our cryo-EM model but was not present in the published crystal model (Fig 4f). In the crystal structure, eight amino acids between F510 and Y519 were absent; in contrast, the MORC2 cryo-EM density map allowed us to build all but one residue. Consequently, we accurately determined the structural context of seven amino acids that were unresolved in the published model.

Collectively, these analyses show that PartiNet enables faster particle picking, improved orientation sampling, and higher-quality reconstructions on previously unseen datasets, facilitating accurate model building and recovery of structural features that were unresolved in prior studies.

### PartiNet enables size-agnostic, single-run identification of multiple species

PartiNet’s size-agnostic detection - which requires no manual specification of particle dimensions - enables simultaneous identification of multiple species in heterogeneous datasets. This eliminates the need for multiple picking runs with different size parameters, a common requirement for existing methods when analyzing samples containing particles of varying sizes. We investigated this capability using two published datasets containing multiple molecular species We first evaluated this on EMPIAR-11043, containing the Ankyrin-1 complex in a micelle, and the free ‘Band 3’ protein (Fig 5)^30^. Processing this dataset proved to be quite involved for the original authors, as identification and processing of the two species required 4 different rounds of particle picking (including manual picking and training a Topaz model) in conjunction with iterative 2D and 3D classification before model refinement. Using PartiNet, we were able to demonstrate a high-speed, single picking step which was able to identify both species using our general model (Fig 5a-b), whilst simultaneously avoiding multiple rounds of particle picking and classification (Supplementary Fig 14). To verify whether both species were picked, we plotted the box size of each PartiNet prediction against the confidence associated with each confidence as a 2D histogram. Bivariate and univariate kernel density estimations identified two distinct populations of box sizes in PartiNet coordinates, corresponding approximately to the expected sizes of Ankyrin-1 and Band 3 (Fig 5c). From these coordinates, we were able to import, extract and refine these particlesubsets, resulting in 3.1 Å and 3.2 Å maps of the Ankyrin-1 complex and Band 3, respectively (Fig 5d-f).

**Figure 5.**
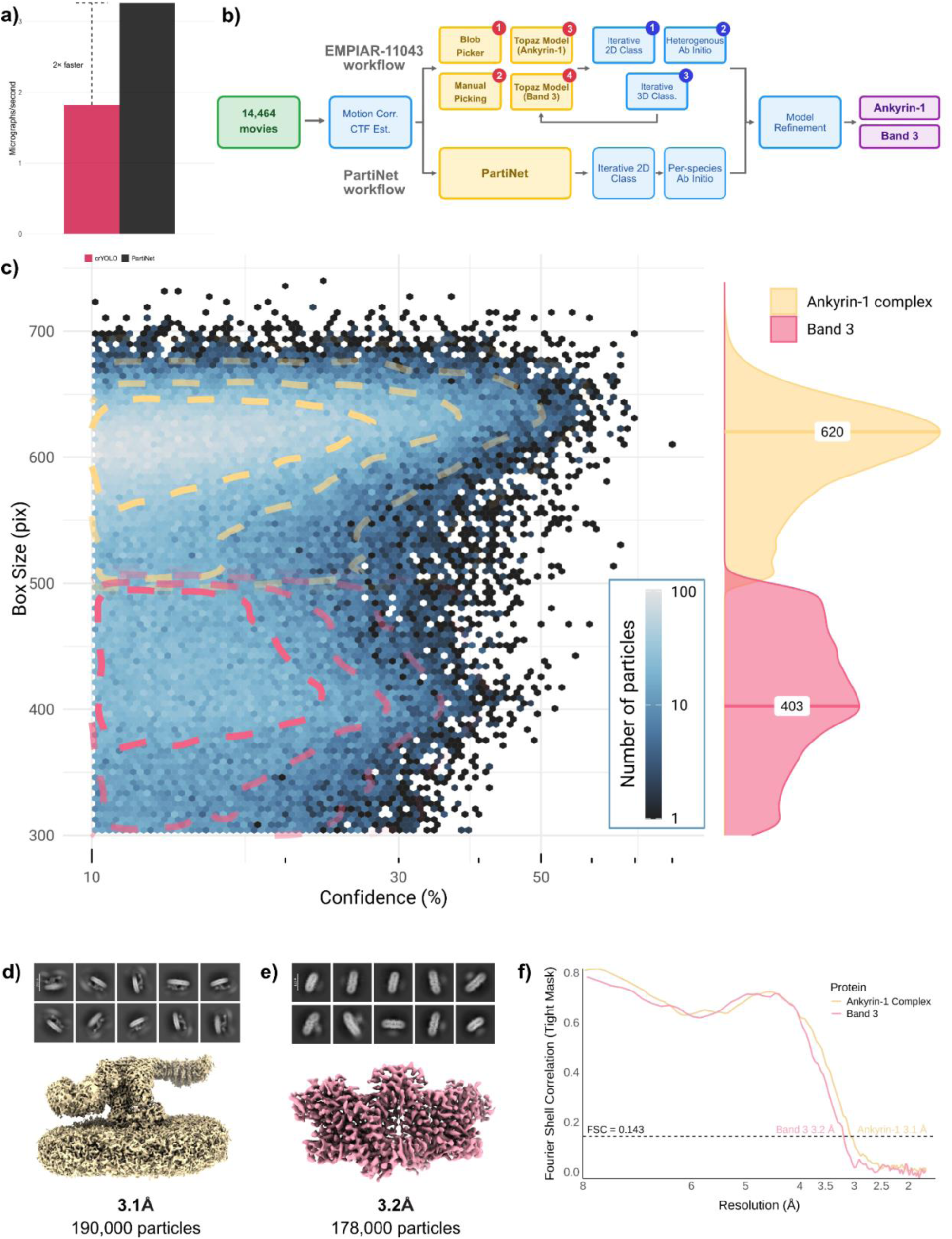
PartiNet can identify multiple species in a dataset. **a.** Bar chart comparing picking speed of PartiNet (black) and crYOLO (red) on EMPIAR-11043. Approximate 2-fold increase in speed with PartiNet is annotated **b.** Simplified workflow for processing EMPIAR-11043 to reconstruct Ankyrin-1 and Band 3 proteins. The published workflow required multiple rounds of particle picking (manual, template-based, and Topaz with trained models) to identify multiple species, whereas PartiNet identifies heterogeneous proteins in a single step. Full workflow details are available in Supplementary Fig 14 **c.** Bivariate histogram showing the relationship between box size and confidence for PartiNet picks on EMPIAR-11043. Particle counts are hexagonally binned with linear box size and logarithmic confidence scaling. Bin color intensity (lighter blue = higher counts) is scaled logarithmically. Only picks with > 10% confidence and > 300 pixel box size are shown to exclude low-quality, noisy detections. Red and yellow contours show bivariate kernel density peaks, with corresponding marginal density plots (right) revealing two distinct box size populations at 620 and 403 pixels, corresponding to Ankyrin-1 complex and Band 3 proteins, respectively. **d-e**. Representative 2D class averages and consensus refinement maps for (d) Ankyrin-1 complex and (e) Free Band 3 proteins identified by PartiNet. Final particle counts for each reconstruction are indicated. **f.** FSC curves for final reconstructions of Ankyrin-1 (yellow) and Band 3 (pink) proteins with resolution annotated at FSC = 0.143 cutoff.

We completed our validation of PartiNet on EMPIAR-12531 containing the MlaCD complex with two species comprising 1:6 and 2:6 stoichiometries of MlaC:MlaD (Fig 6, Supplementary Fig 15)^31^. PartiNet was able to pick 2 times faster than crYOLO on this dataset (Fig 6a). PartiNet also was able to contribute more particles to final reconstruction of each species than the template picking used in the original publication (Fig 6b). PartiNet picked 1,802,428 particles, of which 211,021 were used for 1:6 reconstruction, and 203,014 for 2:6. Conversely, in the original publication, 519,770 particles were picked with template picking and 97,460 particles were used for 1:6 reconstruction and 58,259 for 2:6 reconstruction. Using PartiNet picks, we were able to calculate reconstructions with 1:6 species at 3.7 Å and 2:6 species at 3.4 Å compared to 4.4 Å for each species published previously^31^ (Fig 6c). Smearing of signal was seen in 3D reconstructions of both complexes, suggesting suboptimal “stacking” of the protein complexes in the sample. This necessitated masking for only the complex in final refinements (Supplementary Fig 15). Subsequent reconstructions of both 1:6 and 2:6 species showed well-defined and isotropic secondary structure (Fig 6e-h), with well-resolved interfaces between subunits in the 2:6 complex. Particle distribution plots confirmed distributed sampling across most views for both species, with excellent angular coverage (Fig 6f-i). With the reconstruction of the two species, we were able to confirm the observation from the original authors that the binding of the MlaC subunit to the MlaD multimer breaks the hexameric symmetry of the internal pore, with the MlaD monomer contracting slightly at the centre. We also observed a density in the central pore of the 2:6 species reconstruction; however, we were unable to determine at this resolution if this was the presence of lipid or an artifact from refinement.

**Figure 6.**
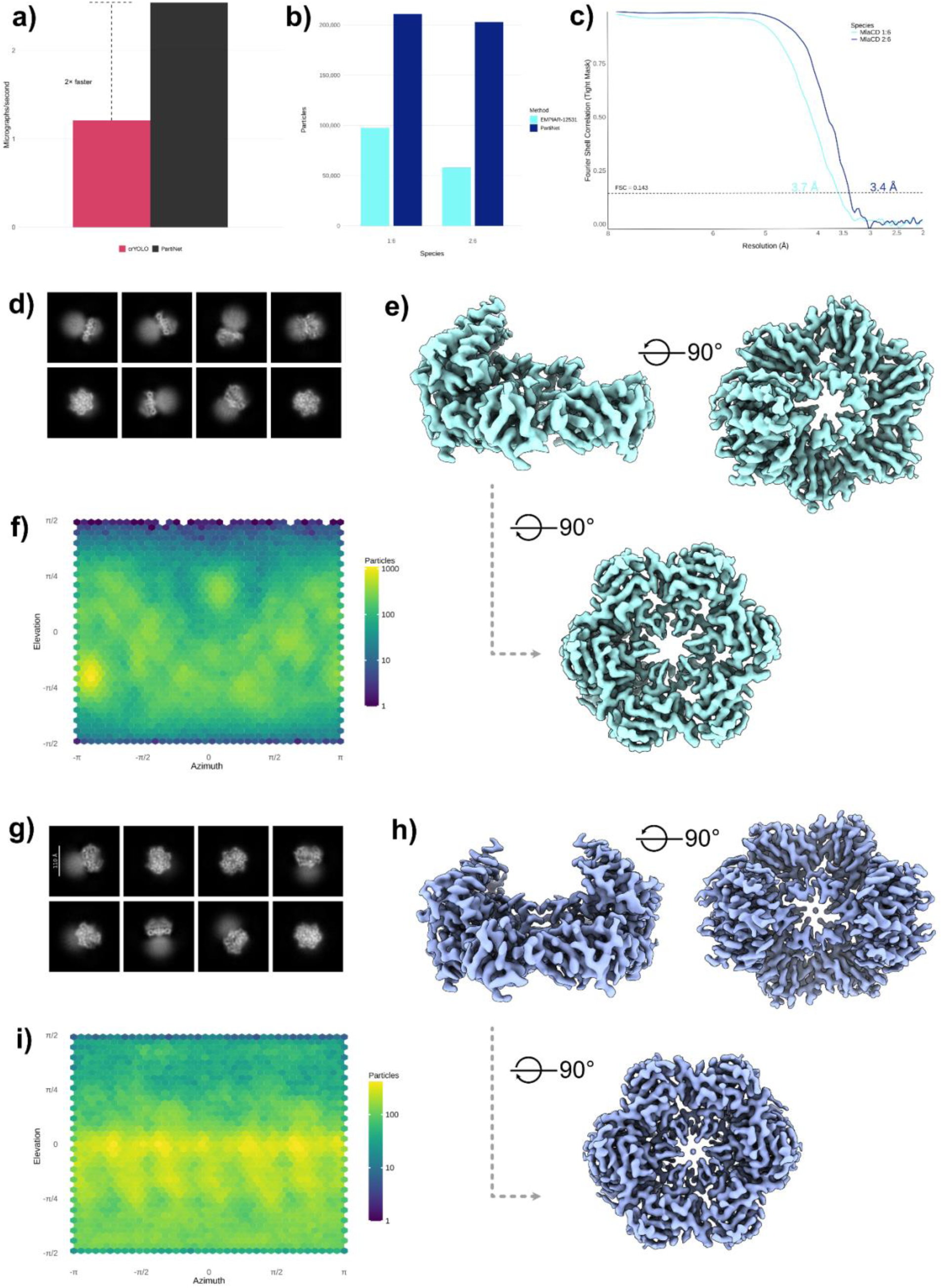
PartiNet improves the resolution of recently published maps. **a.** Bar chart comparing picking speed of PartiNet (black) and crYOLO (red) on EMPIAR-12531. Approximate 2-fold increase in speed with PartiNet is indicated. **b.** Bar chart comparing number of particles used for final reconstruction of the MlaCD complex (both 1:6 and 2:6 stoichiometry species) between published workflow (light blue) and with PartiNet picking (dark blue). **c.** FSC curves for final reconstructions of 1:6 stoichiometry (light blue) and 2:6 stoichiometry (dark blue) species of MlaCD with resolution annotated at FSC = 0.143 cutoff. **d.** Representative 2D class averages of 1:6 stoichiometry MlaCD from particles picked with PartiNet. **e.** Cryo-EM map reconstruction of 1:6 stoichiometry MlaCD from PartiNet particles. **f.** 2D heatmap of particle distributions for 1:6 stoichiometry reconstruction (yellow indicating high particle counts and dark blue low particle counts). **g.** Representative 2D class averages of 2:6 stoichiometry MlaCD from particles picked with PartiNet. **h.** Cryo-EM map reconstruction of 2:6 stoichiometry MlaCD from PartiNet particles. **i.** 2D heatmap of particle distributions for 2:6 stoichiometry reconstruction (yellow indicating high particle counts; dark blue indicating low particle counts).

Taken together, the results from Ankyrin-1 and MlaC/D complexes show that PartiNet can recover multiple molecular species from heterogeneous datasets using a single picking run, improving both throughput and reconstruction quality in multi-component samples.

## Discussion

PartiNet is a dynamic neural network architecture for particle picking in cryo-EM that addresses critical bottlenecks in protein structure determination. By implementing adaptive inference pathways that adjust computational complexity based on micrograph difficulty, PartiNet achieves substantial improvements in both speed and accuracy compared to existing methods. Our comprehensive validation demonstrates consistent improvements in particle identification, reconstruction quality, and computational efficiency across diverse datasets spanning different protein classes, sizes, and imaging conditions.

### PartiNet is a dynamic particle picker for cryo-EM

PartiNet is the first dynamic architecture applied to cryo-EM particle picking. Unlike static networks that process all micrographs through fixed detection layers ^6,12,14–16,18^, PartiNet’s adaptive router adjusts network depth *in situ* based on learned image features. This addresses a fundamental limitation: the inability to modulate computational resources according to image difficulty. Critically, the router learns to identify challenging features *-* complex noise, contamination, variable defocus, support film—without supervision, capturing *imaging* and sample-related complexity without manual labeling. This modularity enables future detector upgrades without architectural redesign and opens possibilities for adaptive approaches in other cryo-EM steps including 3D classification, refinement, model building.

### PartiNet enables truly automated on-the-fly processing workflows

Consistent resolution improvements of up to 1.0 Å suggest PartiNet’s selection criteria better align with reconstruction requirements, reflecting diverse CryoPPP training and dynamic resource allocation for high-quality particle detection. Superior ModelAngelo metrics confirm that improvements translate to more reliable atomic models^37^.

Critically, PartiNet’s combination of speed (2–7× faster) and size-agnostic picking—without prior parameter specification—enables truly automated workflows. Existing methods require manual size input, and they struggle with heterogeneous samples, necessitating user intervention. PartiNet’s YOLO architecture^20^ simultaneously determines presence, location, and dimensions, handling heterogeneous samples (Ankyrin-1/Band 3, MlaCD) in single passes versus multiple rounds with manual intervention.

This transforms cryo-EM practice: real-time particle detection with quality metrics provides immediate assessment of sample quality, grid selection, and defocus optimisation during acquisition rather than days later. Integration with existing on-the-fly platforms - Warp/M^43^, CryoSPARC Live^44^, and RELION^3^ - would enable direct sample assessment during imaging. Microscopists can identify failing samples or optimal parameters without interrupting collection, substantially reducing experimental iteration cycles and microscope time waste, which is particularly valuable for high-throughput facilities processing diverse samples.

### Limitations and considerations for practical deployment

Several limitations warrant consideration. First, extremely unusual morphologies (elongated filaments, <50 kDa proteins, extreme aspect ratios) may benefit from fine-tuning, requiring GPU resources (≥8 GB VRAM) and PyTorch familiarity. Second, speed advantages require multi-GPU setups (our benchmarks: 4× A100 GPUs); single-GPU users may see only modest improvements. Memory requirements (∼64 GB) may limit older hardware. Third, severe micrograph quality issues (motion artifacts, thick ice, damage, contamination) challenge all AI pickers^25^ - no computational approach substitutes for high-quality sample preparation. Fourth, while PartiNet identifies multiple species by size, closely-sized particles with similar contrast require downstream 2D/3D classification. Fifth, PartiNet uses YOLOv7 (2022); newer architectures like YOLOv10^45^ and Detection Transformer^46^ may offer improvements, though the modular framework facilitates future upgrades.

In conclusion, PartiNet demonstrates that dynamic architectures can simultaneously improve speed, accuracy, and robustness in cryo-EM particle picking. Several lessons emerge from this work: diverse training data is essential for broad applicability, modular architectures allow future improvements, combining AI with domain-specific methods (like appropriate denoising) yields better results than AI alone, and open-source distribution accelerates community adoption.

As AI integration into cryo-EM accelerates, approaches balancing automation with flexibility and providing interpretable outputs (confidence scores, difficulty classifications) will be essential. The dynamic architecture paradigm provides a template for future developments that adapt intelligently to biological complexity.

## Online Methods

### Dynamic architecture inference and training

PartiNet is built on a dynamic, deep learning architecture called DynamicDet^24^ and trained on EM micrographs to detect and localise protein particles. Dynamic architectures can adjust the network architectures in response to different imaging and sample conditions, making it possible to train models efficiently, and also to detect particles with increased speed and accuracy^22,23^.

Static object detection algorithms (hereafter called detectors) contain a backbone, neck and head^19,45^. During model inference, the backbone performs the bulk of the feature extraction operations on the input image, the neck pools and aggregates these features, and the head performs the final detection by generating bounding boxes or providing pixel coordinates. This architecture is common across all detector classes. DynamicDet differs from other object detection algorithms because, instead of passing images sequentially through the backbone, neck, and head of a single detector, it evaluates the intermediate outputs at each stage to adjust the network’s size for processing the input image (Fig 1).

Assume for a static Detector *D*_1_ which has backbone *B*_1_, neck and head *H*_1_ such that for a given input image x:

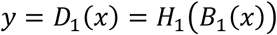

where *y* is the output (coordinates/bounding boxes/contour) of a static object detection network *D*_1_ on input image *x*.

Instead of a static network, DynamicDet performs the following operations on input *x*:

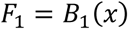

where *F*_1_ are the multiscale feature maps extracted by the first backbone *B*_1_. These feature maps are fed to an adaptive router *R* which determines a difficulty score *ϕ* of the image:

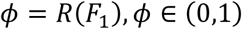

If an image is easy (based on a learned threshold from training, see below) then processing is completed by the first detector:

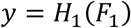

However, if the image is ‘hard’, then these features *F*_1_ are passed along with the input image *x* to the second detector such that:

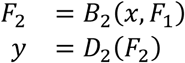

In this way, DynamicDet can dynamically route images to a complex or simpler network based on perceived difficulty of the input images. Currently, PartiNet uses two identical YOLOv7 detector backbones^20^; however there is scope to change these detectors, highlighting the modularity of the DynamicDet framework.

Total training of a model for PartiNet requires two steps: training of the detectors, then training of the adaptive router (Supplementary Fig 16). To train the adaptive router, the model weights of the dual detector are frozen, leaving only the adaptive router to be trained to determine if a micrograph is easy or hard ^24^. This is accomplished with an adversarial loss comparison. A micrograph is passed through a single detector and the total loss ℒ_1_ is calculated. Then, the micrograph is passed through both detectors and total loss ℒ_2_ is calculated. If the difference between the two ℒ_2_ − ℒ_1_ is low, there is no advantage in using two detectors for the micrograph, and the adaptive router learns this micrograph as easy. If ℒ_2_ − ℒ_1_ is high then there is a distinct advantage in using the second detector and the micrograph is labelled as easy. Importantly, this stage is unsupervised: the training dataset does not need to be labelled individually as easy or hard (for example, by a skilled human). Instead, only the micrograph is required, removing bias during training of the adaptive router. The adaptive router is able to encode important optical considerations from the image acquisition into its training regime without the need for explicit labelling. This allows for *in situ* dynamic routing of images based on the difficulty of input micrographs.

### Calculation of Precision, Recall and mAP@50%

In PartiNet, the Intersection over Union (IoU) is used for assessing the overlap between predicted and ground truth bounding boxes. The IoU was calculated as:

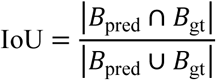

where *B*_pred_ represents the predicted bounding box and *B*_gt_ represents the ground truth bounding box. The confidence score associated with each predicted bounding box represents the model’s confidence of prediction in both the protein particle’s presence and the accuracy of its localisation. PartiNet also calculates Precision and Recall during training:

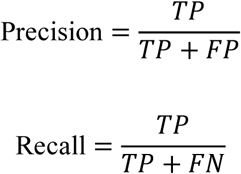

Where:

- *TP* (True Positives): Correctly detected objects with IoU ≥ 0.5
- *FP* (False Positives): Detected objects with no corresponding ground truth
- *FN* (False Negatives): Ground truth objects not detected

The mean Average Precision at 50% IoU threshold (mAP@50%) is calculated as:

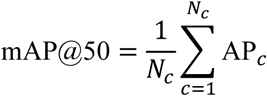

where:

- *N_c_*: Total number of classes (in the case of PartiNet *N_c_* = 1)
- AP*_c_*: Average Precision for each class, computed by integrating the precision-recall curve

The precision-recall curve was generated by varying the confidence threshold and calculating precision and recall at each point. The area under this curve represents the Average Precision for a given class^45^.

Bounding box coordinates are provided with an associated confidence in PartiNet to represent a probabilistic assessment combining a) the probability that protein particle is present in the box, and b) the correct localisation of the box in the micrograph. Confidence in the correct class assignment is only relevant for multi-class inference strategies, which PartiNet does not use. Confidence can be calculated as

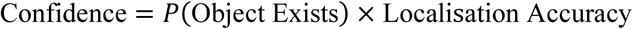

Confidences can be plotted as a histogram to give an initial assessment of the performance of PartiNet. A confidence threshold is specified: bounding boxes with a confidence above this threshold are retained and those below are discarded. PartiNet defaults to a threshold of 0.3 which provides a balance between retaining the majority of particles identified whilst rejecting most junk and spurious picking.

### CryoPPP Image Processing

The CryoPPP test datasets were processed in CryoSPARC v4.6.2 (Supplementary Fig 4). We utilised our trained PartiNet model with a confidence threshold of 0.3, crYOLO’s publicly available model with the “PhosaurusNet” architecture ^15^ at the default confidence threshold of 0.3, and Topaz’s publicly available model with the “ResNet16 (64 units)” integrated directly in CryoSPARC^4,14^ with default parameters to pick particles. We picked particles on all 7 datasets denoised with our integrated denoiser and imported the coordinates alongside the motion-corrected micrographs. We performed CTF estimation on the micrographs to correct for microscope aberrations, then extracted protein particles with an appropriate box size (1.5x largest particle diameter) and performed 2D classification with 50 classes. 2D averages were selected corresponding to protein particles with CryoSPARC’s interactive “Select 2D” function.

These selected particle stacks were used for “*ab initio* Reconstruction”, to coarsely reconstruct an initial 3D map at low resolution without a reference. This coarse reconstruction was then refined using CryoSPARC’s “Homogeneous Refinement” with C1 symmetry for each dataset leading to the final map for evaluation.

### TcdA1 image processing and model building

TcdA1 (EMPIAR-10089) was processed in CryoSPARC v4.6.2. PartiNet, Topaz and crYOLO were compared objectively using a standardised workflow (Supplementary Fig 7). 97 movies were imported, motion-corrected and CTF estimated. Picking was performed using default parameters of each AI picker: the general trained PartiNet model with a confidence threshold of 0.3, crYOLO’s publicly available model with the “PhosaurusNet” architecture^15^at the default confidence threshold of 0.3, and Topaz’s publicly available model with the “ResNet16 (64 units)” integrated directly in CryoSPARC ^4,14^ with default parameters to pick particles. Particles were picked on all 97 micrographs denoised with PartiNet’s integrated denoiser. Particles were extracted with a box size of 384 pixels. Extracted particles were filtered in 3D with multi-class “*ab initio* Reconstruction*”*, and then subjected to iterative “Heterogeneous Refinement” and “Homogeneous Refinement” of the best class using C5 symmetry. After filtering particles in 3D, “Non-Uniform Refinement” was applied with Global CTF Refinement (Tilt, Trefoil, Spherical Aberration, Tetrafoil and Anisotropic Magnification), and optimising per-particle defocus and scale. Reconstruction was completed with “Reference Based Motion Correction” and “Homogeneous Refinement” of motion-corrected particles with C5 symmetry.

Initial model building of TcdA1 was performed with the reconstructed maps from PartiNet, crYOLO and Topaz picked particles using ModelAngelo v1.0.1^37^, an automated AI model-building pipeline (Supplementary Fig 8). Each map along with the FASTA sequence of TcdA1 (UniProt Q9RN43) was input into ModelAngelo and a model prediction was performed. Per-residue confidence scores for each residue were plotted. Root mean square deviation (RMSD) values for a single monomer against the crystal structure of TcdA1 (PDB 4O9Y) were calculated in ChimeraX v1.10.1 using the MatchMaker utility with default parameters and then plotted. The Q-score of each residue was calculated and plotted with ChimeraX using the Q-score plugin^47^.

### Rabbit muscle aldolase image processing and model building

Rabbit muscle aldolase (EMPIAR-10215) was processed in CryoSPARC v4.6.2. PartiNet, Topaz and crYOLO were compared objectively using a standardised workflow (Supplementary Fig 9). 1,021 movies were imported, motion-corrected and CTF estimated. Particles were picked with default parameters for PartiNet, crYOLO and Topaz as outlined previously. Particles were picked on all 1,021 micrographs denoised with PartiNet’s integrated denoiser. Particles were extracted with a box size of 256 pixels. Following extraction of PartiNet particles, “2D Classification” was performed with 50 classes. Due to preferred orientation, the ‘four wing’ view of rabbit muscle aldolase was under-represented. To overcome this, the classes were rebalanced after “Select 2D” with 10 superclasses and a rebalance factor of 0.8 (resulting in ∼ 50% of particles being temporarily discarded), and then initial maps were generated with multiclass “*ab initio* Reconstruction” with C2 symmetry. The initialised maps were inspected and the map which had all four wings present was selected. With this map, “Homogeneous Refinement” with D2 symmetry was performed and included the particles that were discarded during class rebalancing.

Processing was completed with “Non-Uniform Refinement” with Global CTF Refinement (Tilt, Trefoil, Spherical Aberration, Tetrafoil and Anisotropic Magnification), and optimising per-particle defocus and scale with D2 symmetry applied.

Initial model building of rabbit muscle aldolase was performed with ModelAngelo. Each map along with the FASTA sequence of rabbit muscle aldolase (UniProt P00883) was input into ModelAngelo and a model prediction was performed. Per-residue confidence scores for each residue were plotted. The RMSD values for a single monomer against the crystal structure of rabbit muscle aldolase (PDB 6ALD) were calculated in ChimeraX v1.10.1 using the MatchMaker utility with default parameters, and then plotted. The Q-score of each residue was calculated and plotted with ChimeraX using the Q-score plugin^47^ (Supplementary Fig 10).

### Purification of MORC2

MORC2^1-603^ (residues 1-603) was cloned into pFastbac construct with 6xHis at the N-terminus and was purified as described previously^38^. Briefly, MORC2 construct was transformed into EMBacY cells to generate a bacmid. Bacmid DNA was transfected into Sf9 cells using FuGENE transfection reagent (Promega) and virus was passaged twice in the same cell line before large-scale infection. Sf9 cells were a kind gift from the Glukhova lab at WEHI (negative for mycoplasma, identity not independently authenticated by us). For large-scale expression, Sf9 cells at a density of 2–2.5 × 10^6^ cells per mL were infected with 1% (v/v) second passage (P2) virus and incubated at 27 °C for 50–60 h, 220 rpm.

The cells were harvested by centrifugation, resuspended in lysis buffer containing 50 mM HEPES, pH 8.0, 300 mM NaCl, 1 mM TCEP, 5% glycerol, 10 mM imidazole, 10 U per ml benzonase solution (Sigma), 1× cOmplete EDTA-free protease inhibitors (Roche) and 5 mM benzamidine hydrochloride. Cells were lysed by sonication. Clarified cell lysate was incubated with cOmplete His-tag purification resin (Merck) for 1 h followed by wash with lysis buffer. MORC2 proteins were eluted in elution buffer (50 mM HEPES pH 8.0, 300 mM NaCl, 1 mM TCEP, 5% glycerol and 250 mM imidazole). Further purification was performed via size exclusion chromatography using the Superose 6 Increase 10/300 GL column (Cytiva) in SEC buffer (50 mM HEPES pH 8.0, 300 mM NaCl, 1 mM TCEP).

### Surface Plasmon Resonance (SPR)

SPR binding studies of MORC2^1-603^ to H3K9me3 were performed using a Biacore S200 Instrument (Cytiva). Biotinylated H3K9me3 peptide (Active Motif) was diluted to 5 µg/mL in SPR running buffer (10 mM HEPES pH 7.4, 300 mM NaCl, 3 mM EDTA and 0.05% (v/v) Tween-20) to a final immobilisation level of 200-220 response units (RU) on the Streptavidin chip (Cytiva). A blank activation/deactivation was used for the reference surface. DNA binding studies were performed at 20 °C in SPR running buffer. MORC2 proteins were diluted to 1000 nM stock in SPR running buffer and prepared as an 8-point concentration series (2-fold serial dilution, 7-500 nM). Samples were injected in a multi-cycle run (flow rate 30 µL/min, contact time of 60 s, dissociation 120 s) with regeneration with 0.5 M EDTA buffer. Sensorgrams were double referenced, and steady-state binding data were fitted using a 1:1 binding model using Biacore S200 Evaluation Software (Cytiva, v1.1). Representative sensorgrams and fitted dissociation constant (*K*_D_) values are depicted as mean ± SEM (n=3 independent experiments).

### MORC2 cryo-EM sample preparation and data collection

A 0.5 mg/mL MORC2 (residues 1-603) protein solution was incubated with 1 mg/mL biotinylated H3K9me3 peptide (Active Motif, Catalogue number 81047, peptide sequence = ARTKQTAR-Kme3-STGGKAPRKQLA - GGYK (Biotin) - NH2) and 2.5 mM AMP-PNP for 1 h on ice. The MORC2-H3K9me3 sample was applied to UltrAuFoil R1.2/1.3 gold grids (Quantifoil). The grids were glow-discharged for 120 s before application of 4 µL sample onto the grid. The sample was subsequently blotted for 3.5 s (Blot force -3) and vitrified by plunging into liquid ethane using a Vitrobot Mark IV (ThermoFisher) operated at 4 °C and 100% humidity. Cryo-EM data were acquired on a Titan Krios transmission electron microscope (ThermoFisher) operated at 300 keV, equipped with a K3 direct electron detector (Gatan, Pleasanton) with a pixel size of 0.82 Å/pixel and electron dose of 60 e/Å^2^.

### MORC2 Image Processing and model building

MORC2 data was processed in CryoSPARC v4.6.2. PartiNet, Topaz and crYOLO were compared using a standardised workflow (Supplementary Fig 12). 6,148 movies were imported, motion corrected and CTF estimated. Particles were picked with default parameters for PartiNet, crYOLO and Topaz as outlined previously. Particles were picked on all 6,148 micrographs denoised with PartiNet’s integrated denoiser. Particles were extracted with a box size of 256 pixels. Classes were selected from a single round of 2D classification. Maps were initialised for each with “*ab-initio* Reconstruction”, and “Homogeneous Refinement” with C2 symmetry was performed. A final round of “Non-Uniform Refinement” with Global CTF Refinement (Tilt, Trefoil, Spherical Aberration, Tetrafoil and Anisotropic Magnification), and optimising per-particle defocus and scale with D2 symmetry was applied.

### Ankyrin-1 and Band 3 Image Processing

EMPIAR-11043 was processed in CryoSPARC v4.7.0 (Supplementary Fig 14). 14,926 movies were imported, motion-corrected and CTF estimated. Particles were picked with PartiNet on all 14,926 micrographs denoised with the integrated denoiser. 3,614,613 particle coordinates were imported into CryoSPARC, and particles were extracted with a box size of 600 pixels, Fourier downsampled to 150 pixels. Particle stacks were cleaned by 2 rounds of 2D classification (each with 100 classes), with classes selected showing clear averages of both species. Another 2 rounds of 2D classification with 50 classes were generated, with classes selected showing the Ankryin-1 complex in micelle, resulting in 190,334 particles. Ankyrin-1 particles were then re-extracted with a box size of 600 pixels with aligned shifts from 2D. The full resolution particles were initialised with single class “*ab initio* Reconstruction” and then “Homogenous Refinement” in C1. Finally, “Non-Uniform Refinement” was applied with global CTF correction (Tilt and Trefoil) and minimising over per-particle defocus and scale.

In a parallel processing pathway, 2 rounds of 2D classification with 50 classes were done and classes showing Band 3 protein were identified, resulting in 177,900 particles. Band 3 particles were then re-extracted with a box size of 320 pixels with aligned shifts from 2D. The full resolution particles were initialised with single class “*ab initio* Reconstruction” and then “Homogenous Refinement” in C2. To complete processing of Band 3, “Non-Uniform Refinement” was applied with global CTF correction (Tilt, Trefoil, Spherical Aberration, Tetrafoil, Anisotropic Magnification) and minimising over per-particle defocus and scale with C2 symmetry.

### MlaCD Image Processing

EMPIAR-12531 was processed in CryoSPARC v4.7.0 (Supplementary Fig 14). 9,046 movies were imported, motion-corrected and CTF estimated. Particles were picked with PartiNet on all 9,046 micrographs denoised with the integrated denoiser. 1,802,428 particle coordinates were imported into CryoSPARC, and particles were extracted with a box size of 350 pixels and Fourier downsampled to 144 pixels. Two rounds of 2D classification with 150 classes were used to filter junk particles, resulting in 1,535,431 particles. These particles were initialised with “*ab initio* Reconstruction” with 4 classes.

The classes were then heterogeneously refined. The two classes representing the 1:6 and 2:6 were selected for “Non-Uniform Refinement” with C1 and C2 symmetry applied, respectively. The “Heterogeneous Refinement” and “Non-Uniform Refinement” step was repeated 10 times, to effectively filter particles in 3D. The particles corresponding to each 3D class of 1:6 and 2:6 species were then extracted independently at 350 pixels, with updated 2D and 3D aligned shifts. For the 1:6 species, particles were reconstructed with alignments and then “Non-Uniform Refinement” was applied with C1 symmetry, minimising over particle scale. The particles were then corrected for global CTF aberrations (Tilt, Trefoil, Tetrafoil, Spherical Aberration and Anisotropic Magnification) over 3 iterations. The CTF corrected particles were then used for another “Non-Uniform Refinement” with C1 symmetry. Because some particles were stacked on top of each other in 2D projections, spurious density was observed outside the main refinement of the complex, affecting particle alignment and FSC. A mask was generated with “Volume Tools”, with a lowpass filter = 10 Å, threshold = 0.104, dilation radius = 20 pixels, soft-padding = 25 pixels. This mask was used for a final “Local Refinement”.

In a parallel processing pathway, the 2:6 species particles were reconstructed with alignments as above and then “Non-Uniform Refinement” was applied with C2 symmetry, minimising over particle scale. The particles were then corrected for global CTF aberrations (Tilt, Trefoil, Tetrafoil, Spherical Aberration and Anisotropic Magnification) over 2 iterations. The CTF-corrected particles were then used for another “Non-Uniform Refinement” with C2 symmetry and optimising per-particle defocus. Again, the spurious density from suboptimal particle stacking was observed. A mask was generated with “Volume Tools”, with a lowpass filter = 10 Å, threshold = 0.121, dilation radius = 20 pixels, soft-padding = 24 pixels. This mask was used for a final “Local Refinement” with C2 symmetry and optimising per-particle defocus.

### Map visualisation and data plotting

Visualisation of EM maps and atomic coordinates for analysis and figures were done in UCSF ChimeraX v1.10^48^. Charts for analysis and figures were generated in RStudio v 2025.09.0 with R v4.5.1. All figure layouts and exports were done in Affinity Designer v2.6.0

## Data availability

CryoEM maps have been deposited in the EM Data Bank with the following accession codes: EMD-68600 (MORC2^H3K9-bound^). Atomic coordinates have been deposited in the Protein Data Bank with the accession codes 22QM (MORC2^H3K9-bound^). The raw micrographs, particle coordinates from PartiNet, Topaz and crYOLO for MORC2^H3K9-bound^ have been deposited in EMPIAR the following accession codes (EMPIAR-13226). All other cryo-EM maps for other datasets are accessible from https://doi.org/10.57967/hf/7618.

## Code availability

Source code for PartiNet is publicly available on GitHub at https://github.com/WEHI-ResearchComputing/PartiNet. PartiNet is licensed under the MIT License. The model weights can be found here: https://huggingface.co/MihinP/PartiNet. The documentation can be found here: https://wehi-researchcomputing.github.io/PartiNet/.

## Acknowledgements

We acknowledge use of transmission electron microscopes at the Monash University Ramaciotti Centre for Cryo-Electron Microscopy and Ian Homes Imaging Centre, Bio21. We thank the WEHI Cryo-EM Facility, the WEHI Research Computing Platform and Milton high-performance computing facility, the University of Melbourne Spartan high-performance computing facility and the Monash University MASSIVE high-performance computing facility for providing facilities and support. We thank Nicholas Kirk, Alisa Glukhova and Peter Czabotar for their comments on the manuscript. This work was initially supported by WEHI’s New Medicines and Advance Technology funds to AL and SS. MP is supported by Research Training Program (RTP) Scholarship from Faculty of Engineering and Technology, University of Melbourne and Graeme Clark Institute for Medical Engineering Top-Up scholarship. WT is supported by an NHMRC Investigator Grant (GNT 2026635). JDB received support from an Australian Research Council Future Fellowship (FT220100319) funded by the Australian Government. AL is supported in part by funds from the estate of Akos and Marjorie Talon. SS is supported by funds from WEHI, the estate of Akos and Marjorie Talon, The University of Melbourne Attraction and Retention Funds, the NHMRC Investigator grant (GNT 2016827), the Australian Research Council Discovery Project grant (DP250100450), the US Department of Defence Rare Cancer Research Concept Award (HT9425-24-1-0922) and the US Department of Defence Lung Cancer Research Program Award (HT94252510699).

## Author contributions

MP, OJ, MA wrote the initial scripts for PartiNet, with MP developing the final version. WT purified MORC2, prepared cryoEM samples and performed SPR experiments. HV collected cryoEM data. EY contributed to program parallelisation, debugging, and packaging of PartiNet. JI prepared training data. MP and SS conceived the project. JI, JDB, AL and SS supervised the project. AL and SS acquired the funding. MP, JDB, AL and SS analysed the data and wrote the manuscript with contributions from all authors.

