## Supplementary Information for "PartiNet is a dynamic adaptive neural network for high-performance particle picking in cryo-electron microscopy"

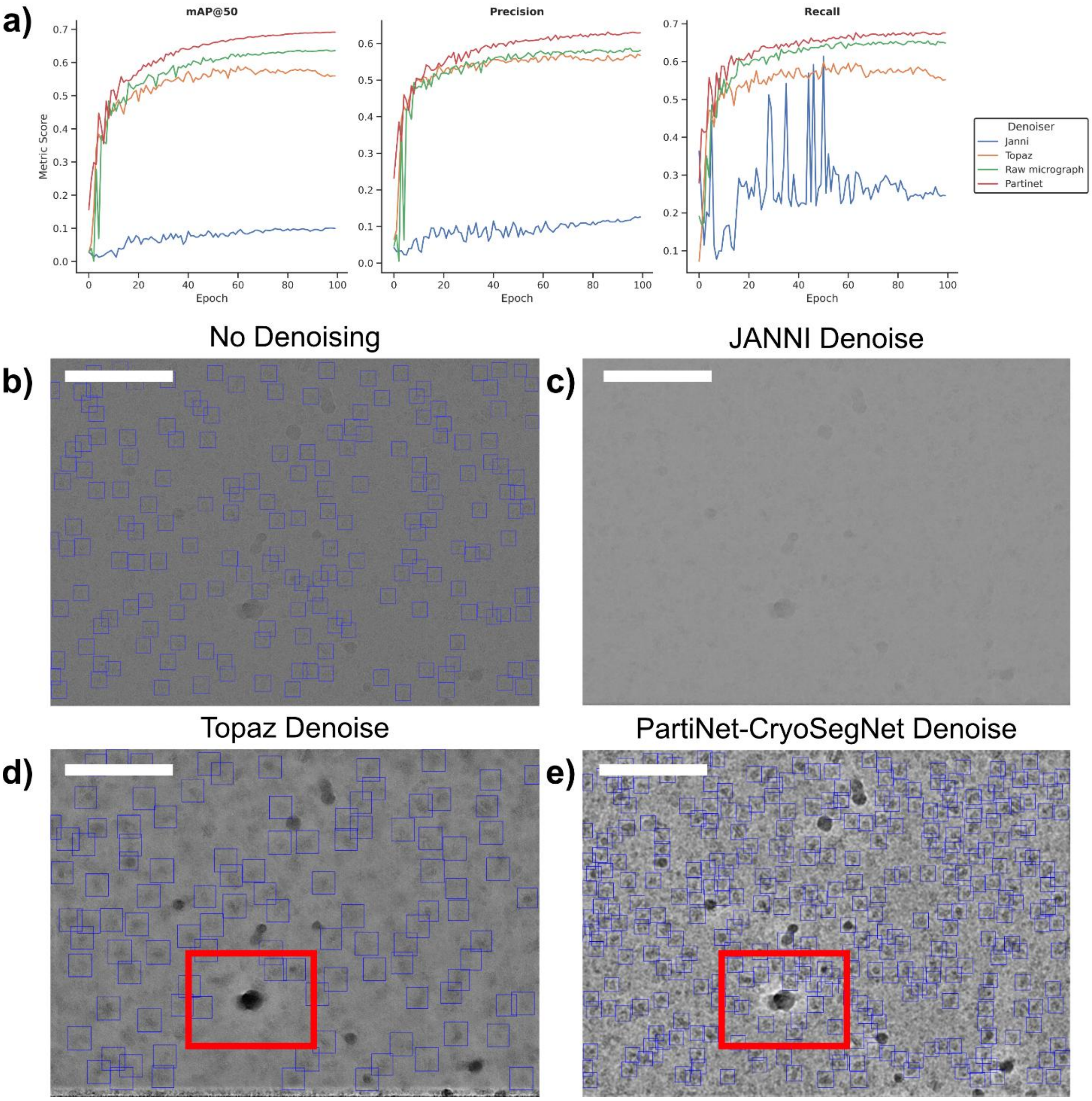

a)
mAP@50
Precision
Recall
Metric Score
Epoch
Denoiser
Janni
Topaz
Raw micrograph
Partinet
No Denoising
JANNI Denoise
b)
c)
Topaz Denoise
PartiNet-CryoSegNet Denoise
d)
e)

### Supplementary Figure 1.

**Comparison of popular denoisers on a micrograph from EMPIAR-10096.**

**a.** Comparison of PartiNet mAP@50%, Precision and Recall during 100 epochs of training on the CryoPPP dataset with raw motion-corrected, JANNI, Topaz or PartiNet-CryoSegNet denoised micrographs. **b.** Raw motion corrected micrograph, PartiNet is able to identify the most obvious particles. **c.** JANNI denoised micrograph. JANNI has effectively flattened the micrograph, and PartiNet was not able to identify any particles in the micrograph. **d.** Topaz denoised micrograph. The presence of the signal dense contamination (identified with the red box) has caused Topaz to flatten all signal around it. Very few particles are picked around this contamination. **e.** PartiNet-CryoSegNet denoised micrograph. The micrograph is effectively denoised with protein particles contrast enhanced, allowing for effectively delineation of closely packed particles, especially around contamination. Scale bar is 80nm.

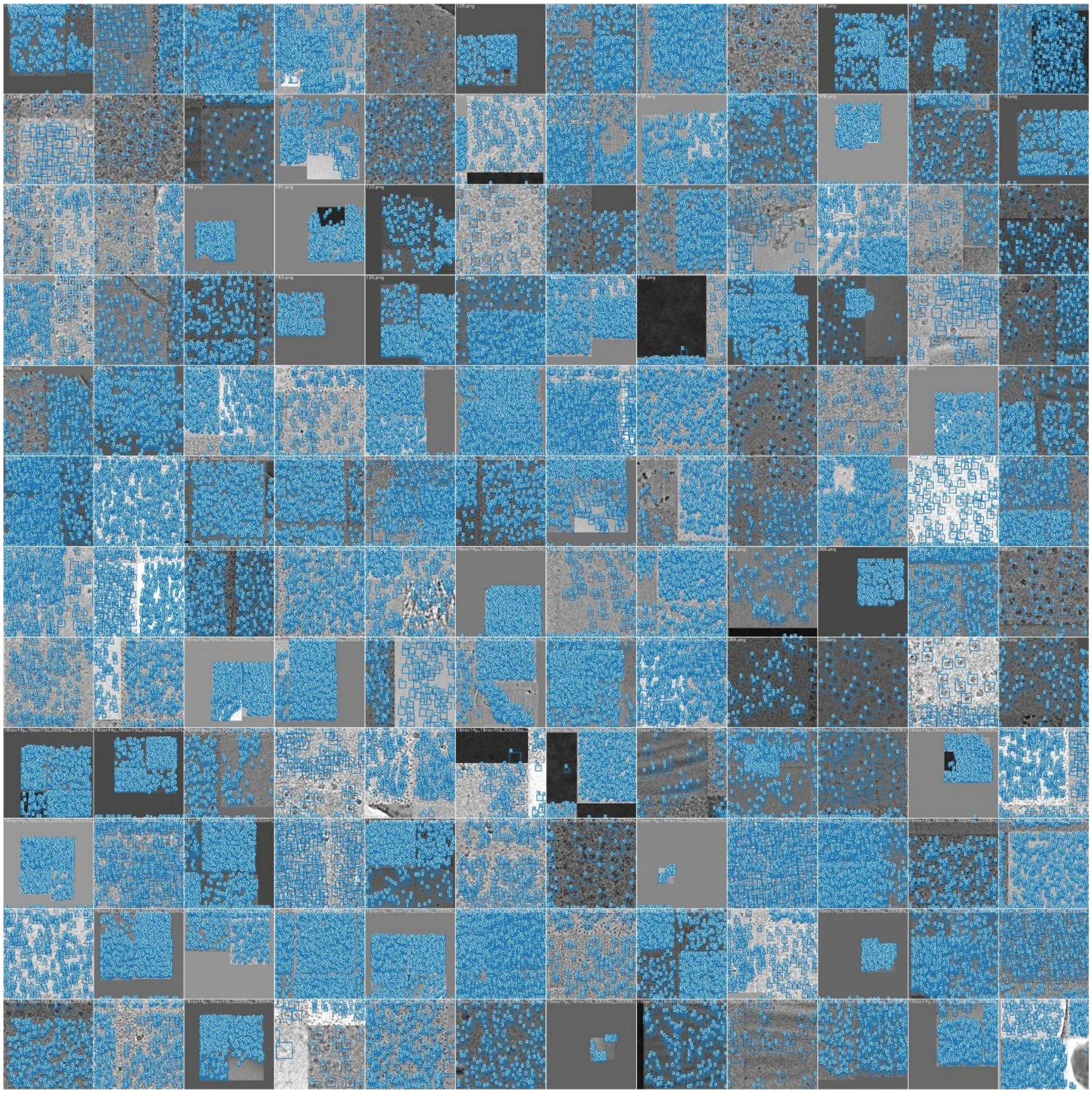

### Supplementary Figure 2.

**Mosaic of image augmentations during PartiNet training effectively increase the feature space of the training data without requiring additional micrographs**

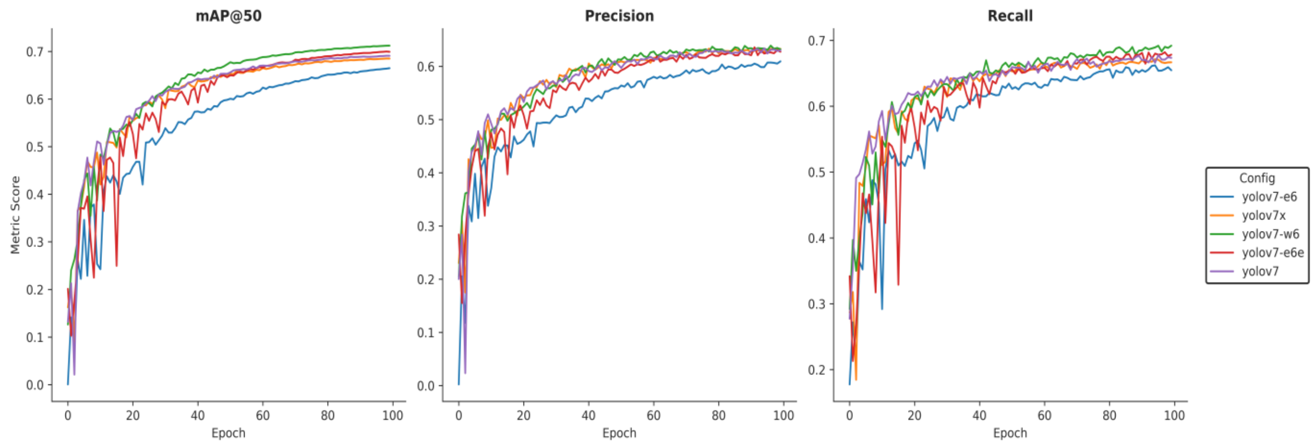


**Supplementary Figure 3.**

**Comparison of different YOLOv7 configurations.**

YOLOv7-W6 was able to slightly outperform other configurations across mAP@50, Precision and Recall. Measured performance of each detector on electron micrographs did not match the reported performance by the original authors of YOLOv7 due to complex characteristics of micrographs. YOLOv7-W6 was used for all subsequent testing and analysis.

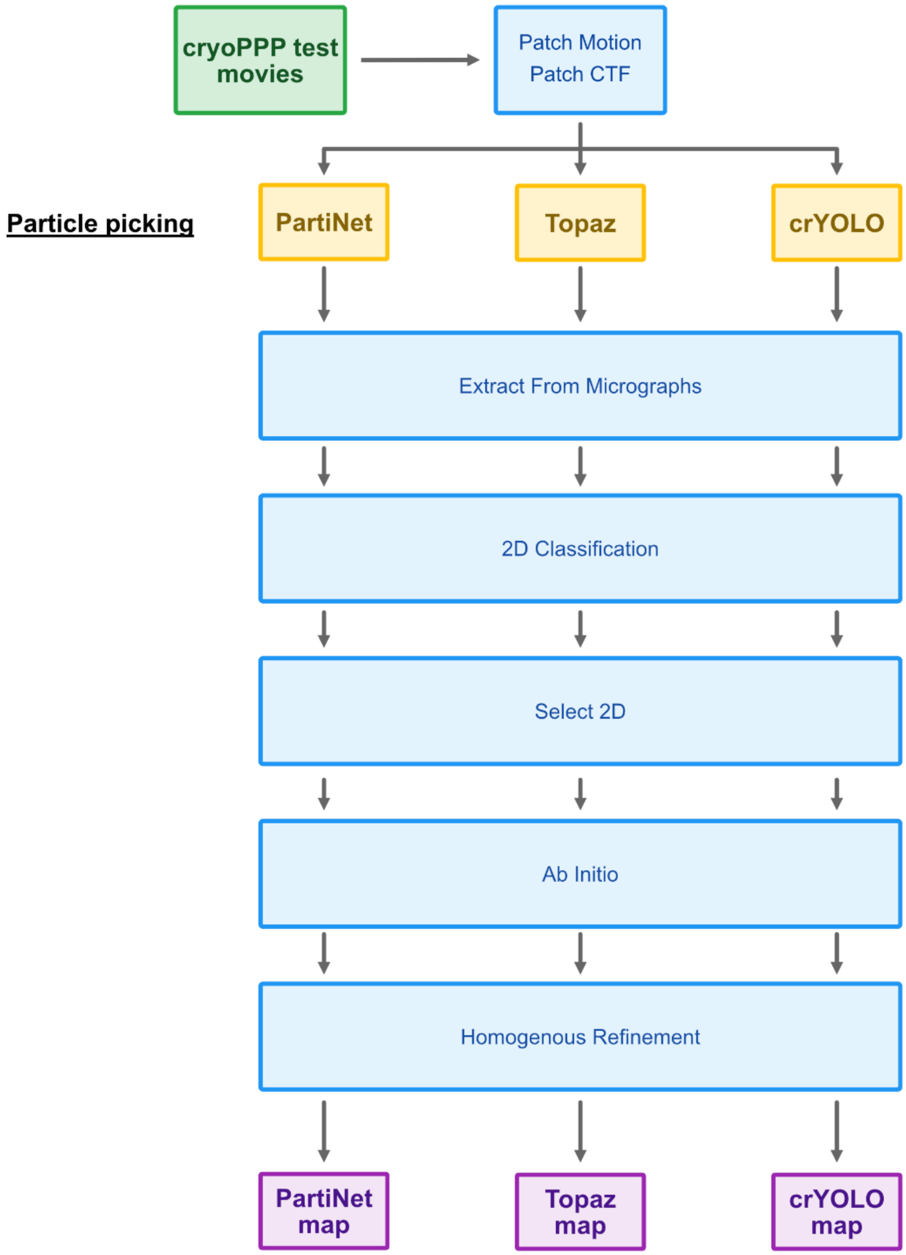


**Supplementary Figure 4.**

**Workflow for cryo-EM processing of CryoPPP test sets.**

Details for processing each test set from CryoPPP can be found in Methods. Processing was done in CryoSPARC v4.6.2

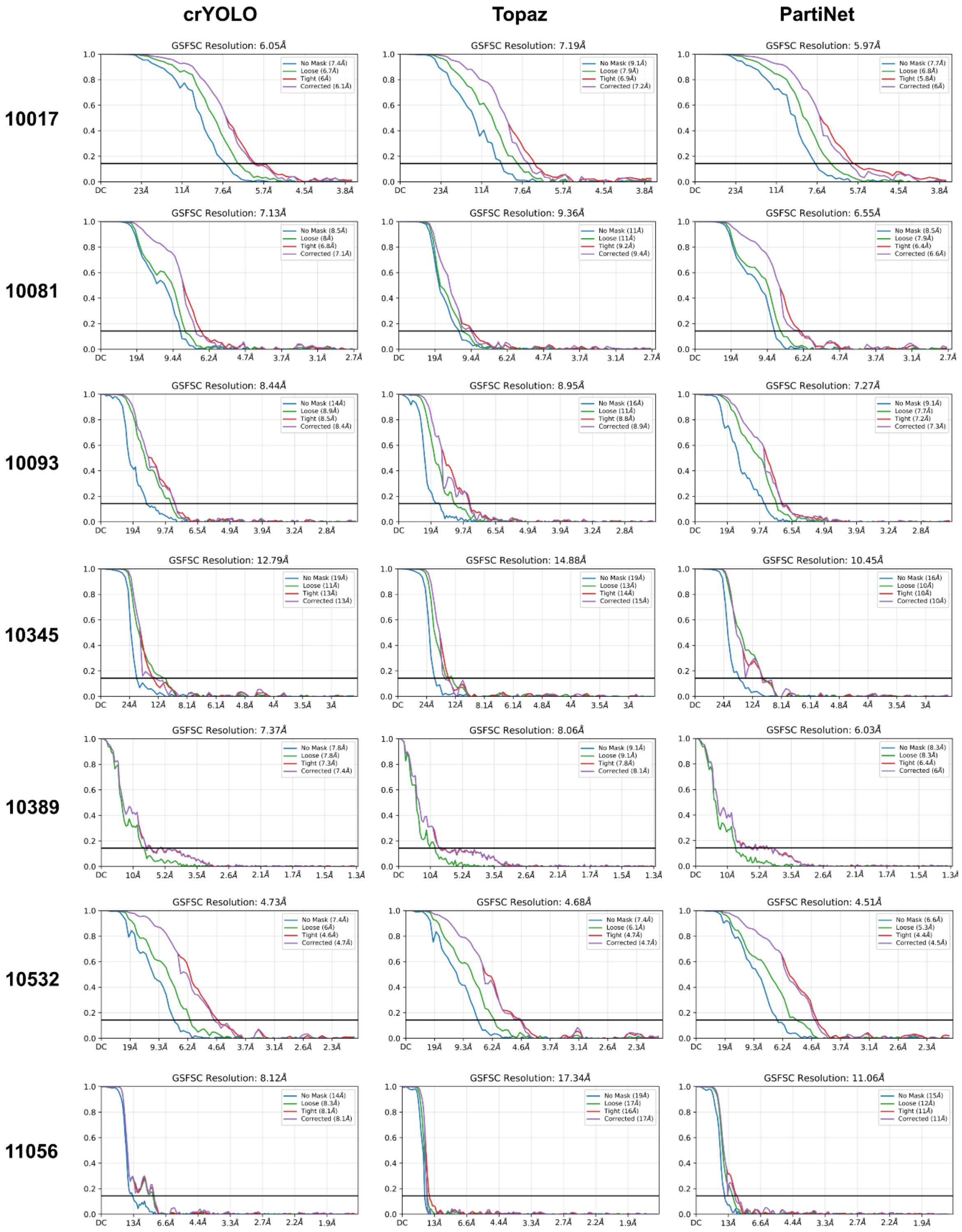


### Supplementary Figure 5.

**FSC curves for CryoPPP test sets.**

Plots were generated in CryoSPARC v4.6.2

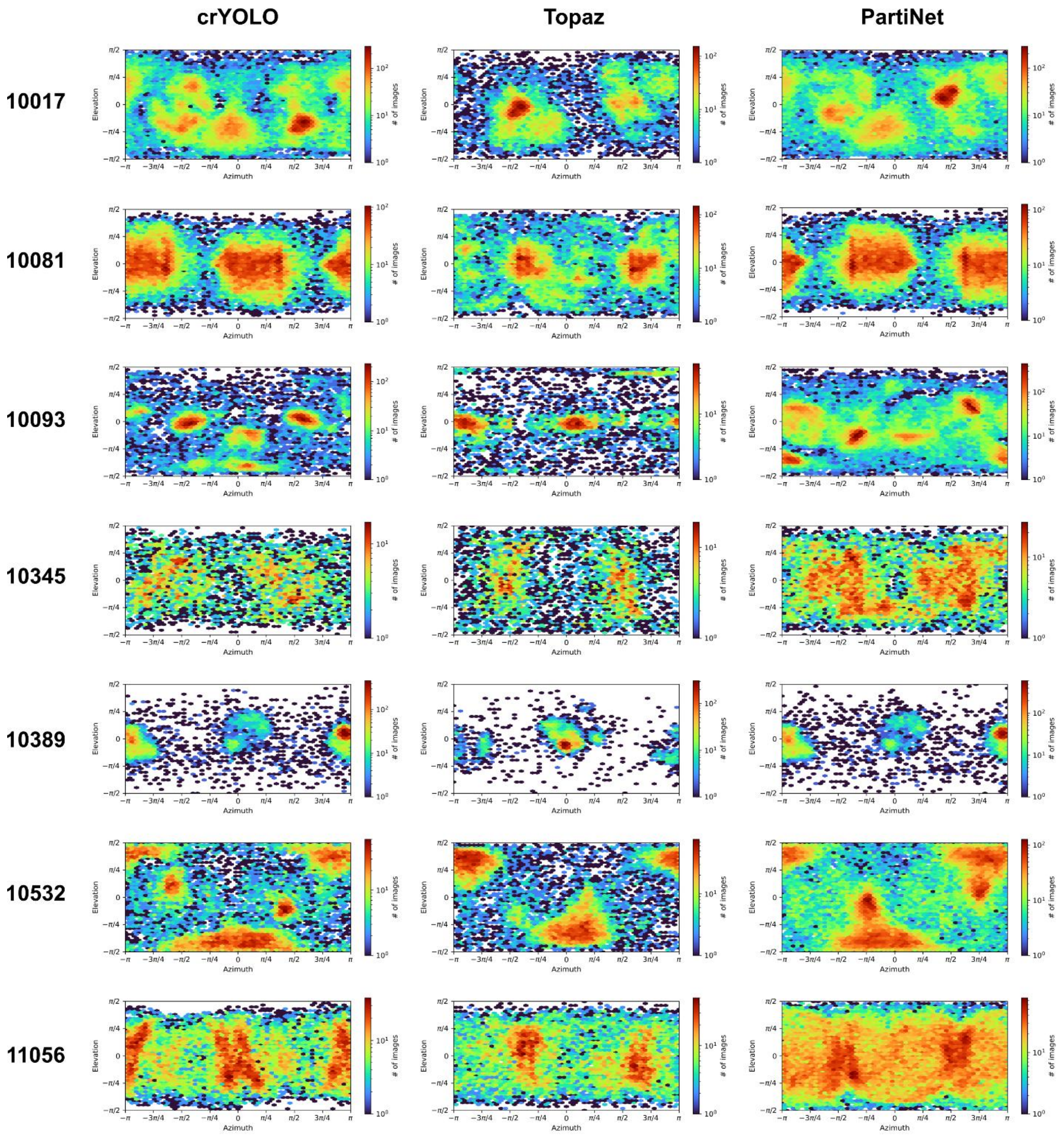


### Supplementary Figure 6.

**Euler maps for CryoPPP test sets**

Plots were generated in CryoSPARC v4.6.2

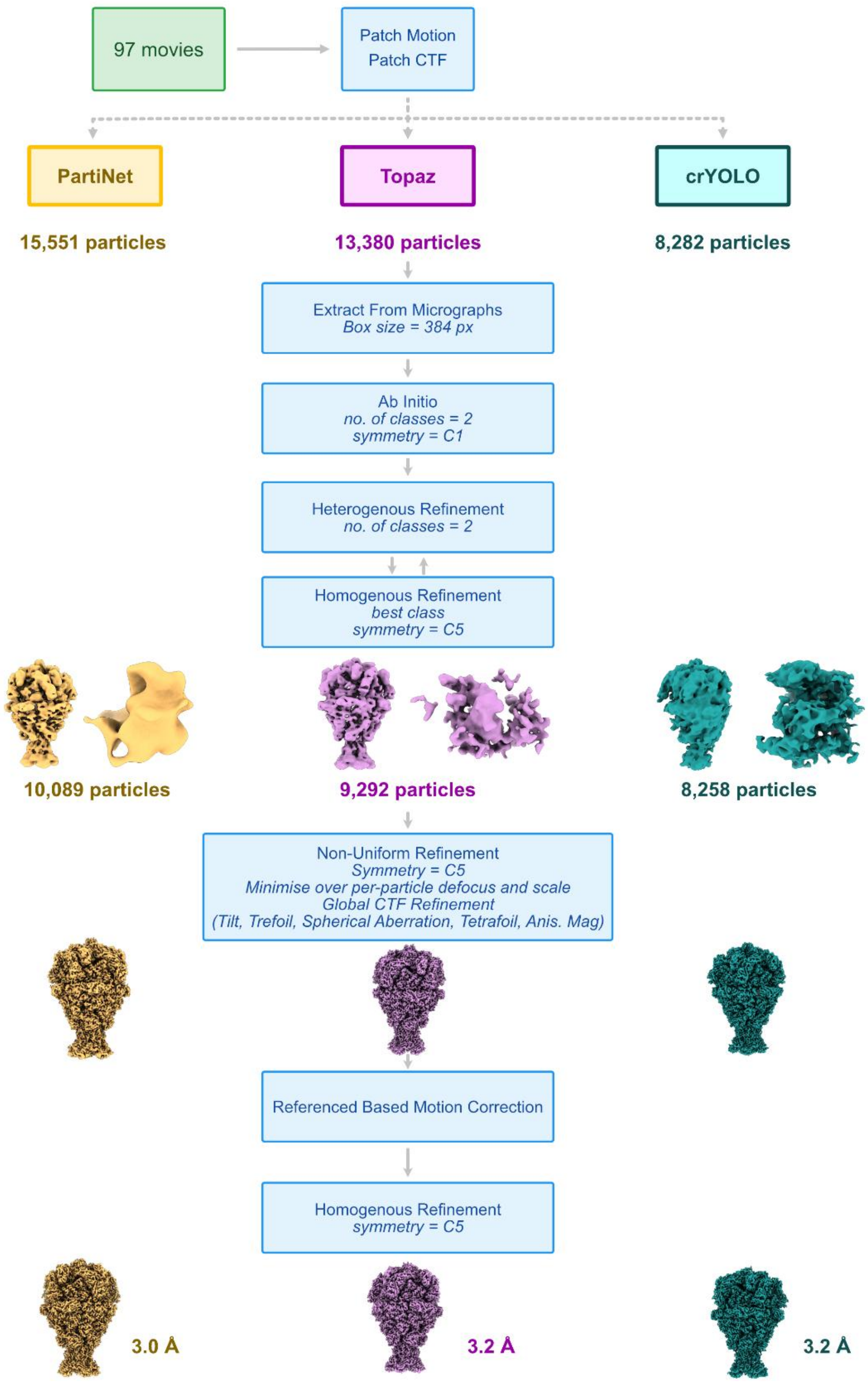


**Supplementary Figure 7.**

**Workflow for processing EMPIAR-10089 (TcdA1)**

Details can be found in Methods. Processing was done in CryoSPARC v4.6.2

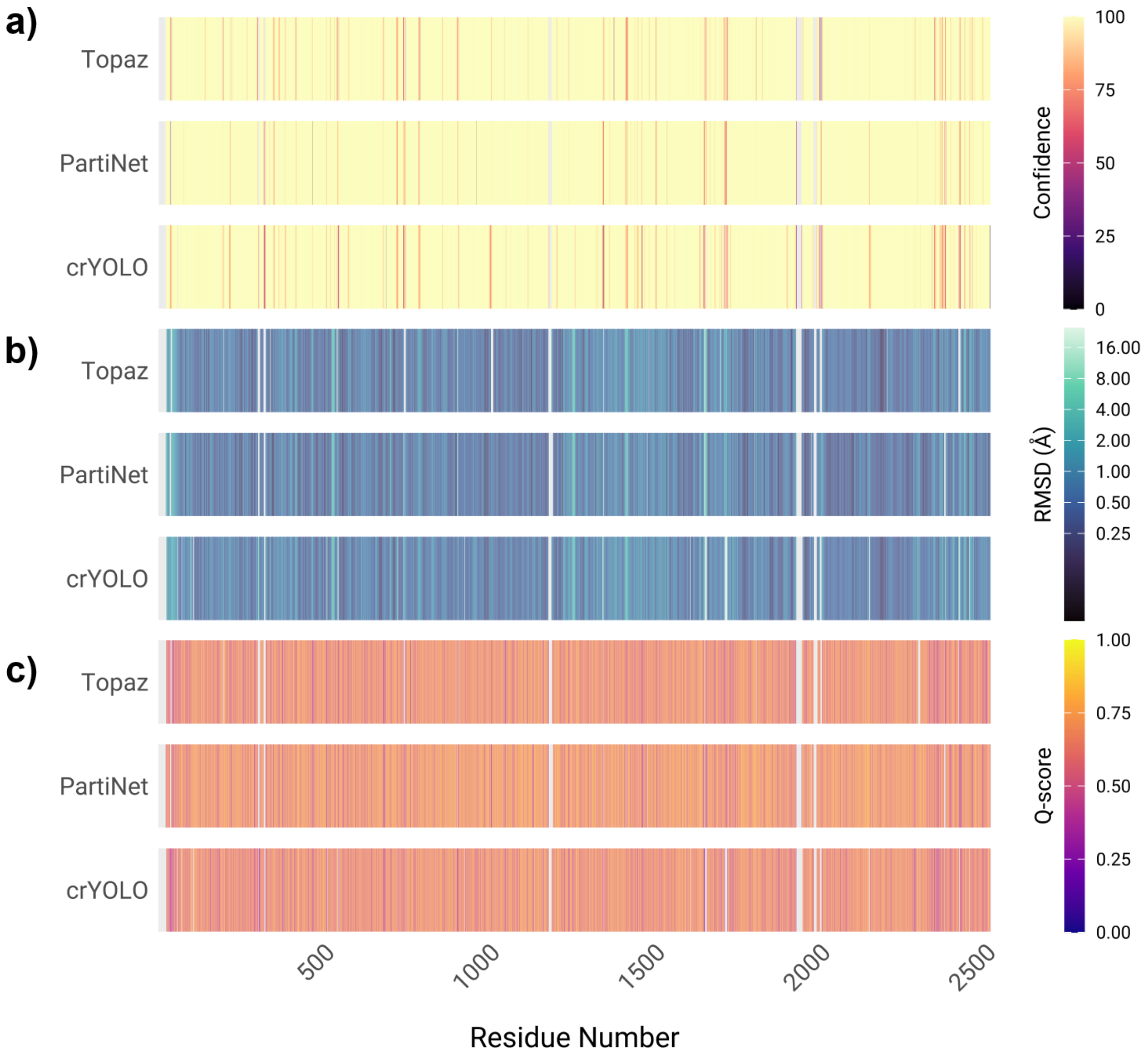


**Supplementary Figure 8.**

**ModelAngelo results for TcdA1 reconstruction.**

**a.** Per-residue confidence of ModelAngelo predictions for TcdA1 maps reconstructed with Topaz, PartiNet and crYOLO. **b.** Per-residue RMSD between ModelAngelo prediction and crystal structure of TcdA1 (PDB 4O9Y). **c.** Per-residue Q-score of ModelAngelo prediction and maps.

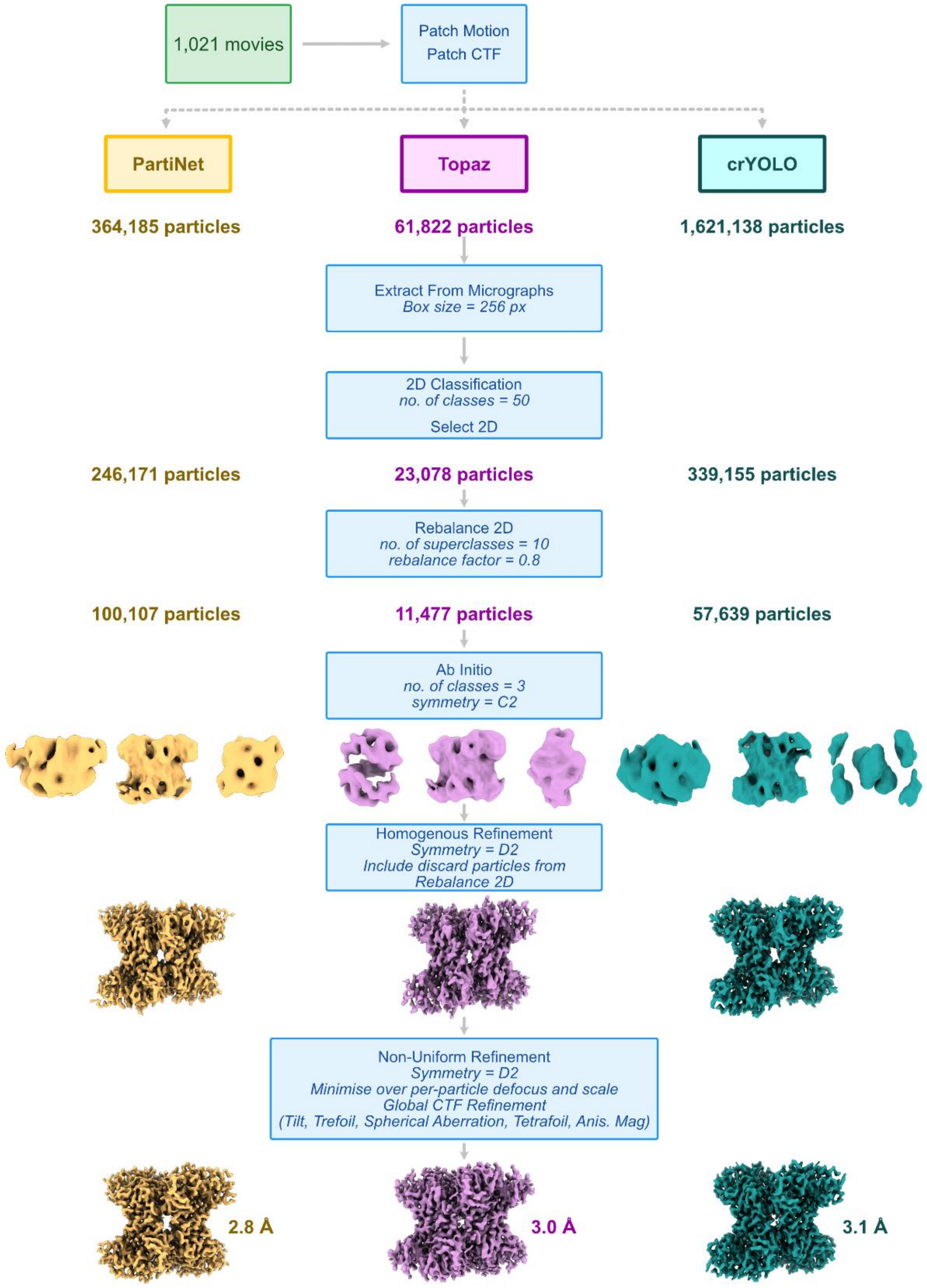


### Supplementary Figure 9.

**Workflow for processing EMPIAR-10215 (rabbit muscle aldolase)**

Details can be found in Methods. Processing was done in CryoSPARC v4.6.2

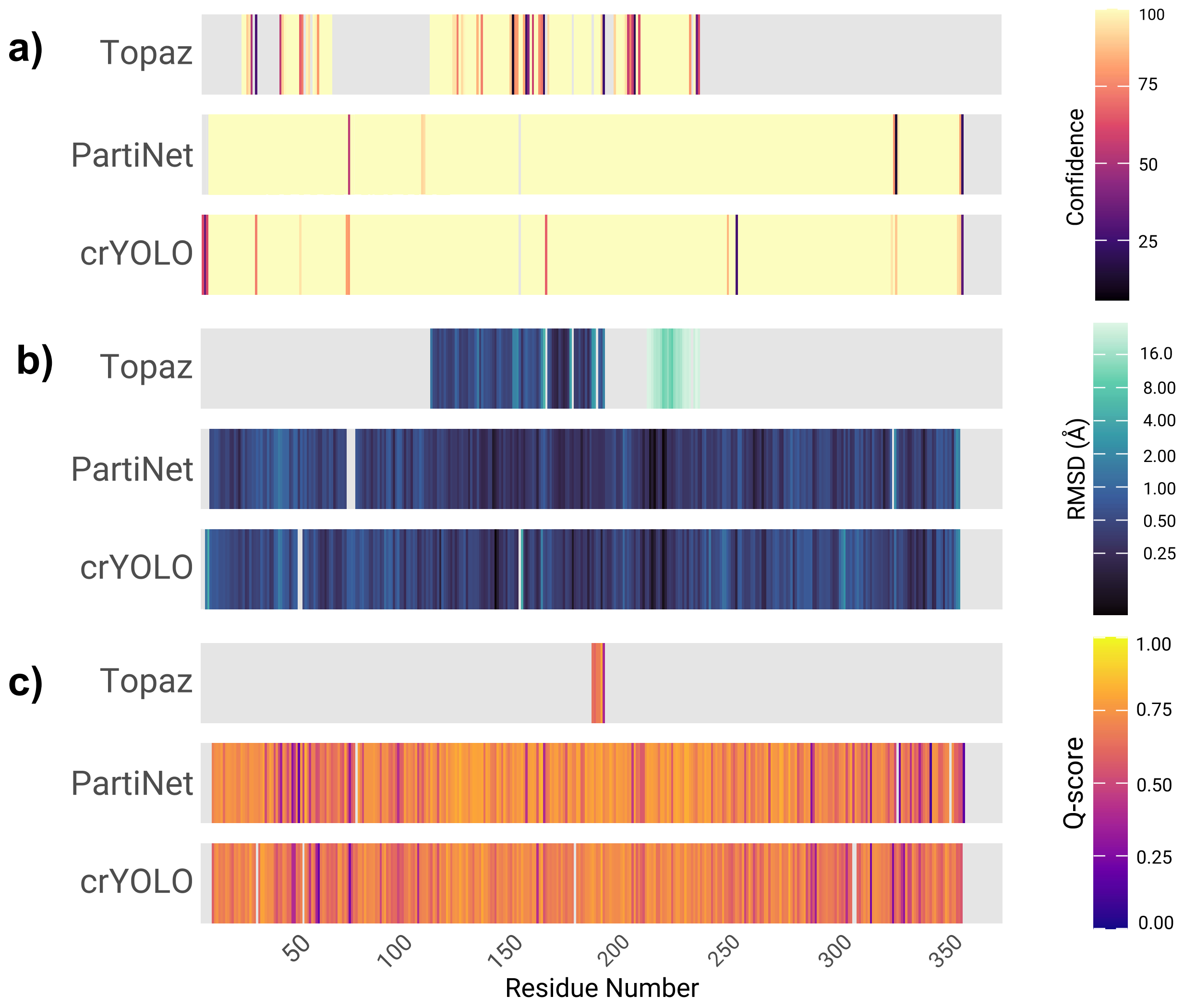


**Supplementary Figure 10.**

**ModelAngelo results for EMPIAR-10215 (rabbit muscle aldolase)**

**a.** Per-residue confidence of ModelAngelo predictions for rabbit muscle aldolase maps reconstructed with Topaz, PartiNet and crYOLO. **b.** Per-residue RMSD between ModelAngelo prediction and crystal structure of rabbit muscle aldolase (PDB 6ALD). **c.** Per-residue Q-score of ModelAngelo prediction and maps.

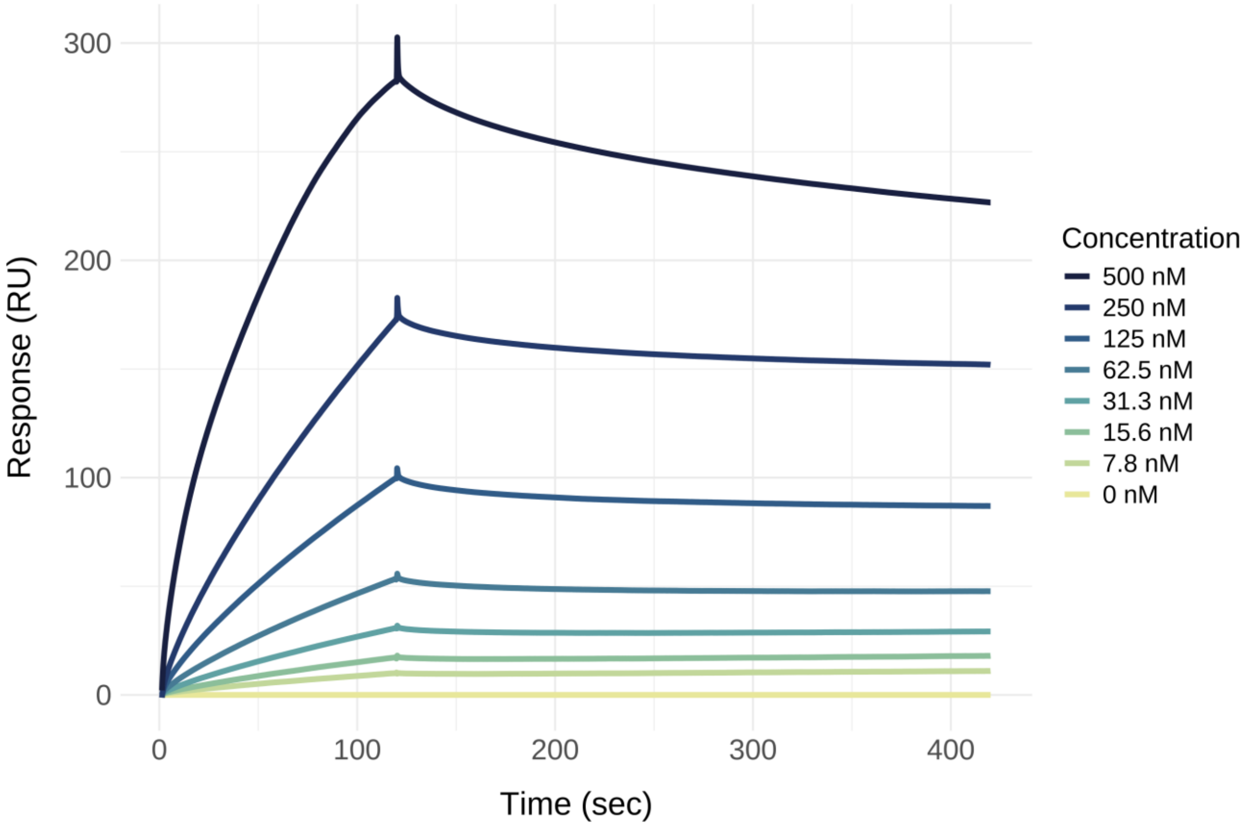


**Supplementary Figure 11.**

**Surface plasmon resonance analysis of MORC2 binding to H3K9me3 peptide.**

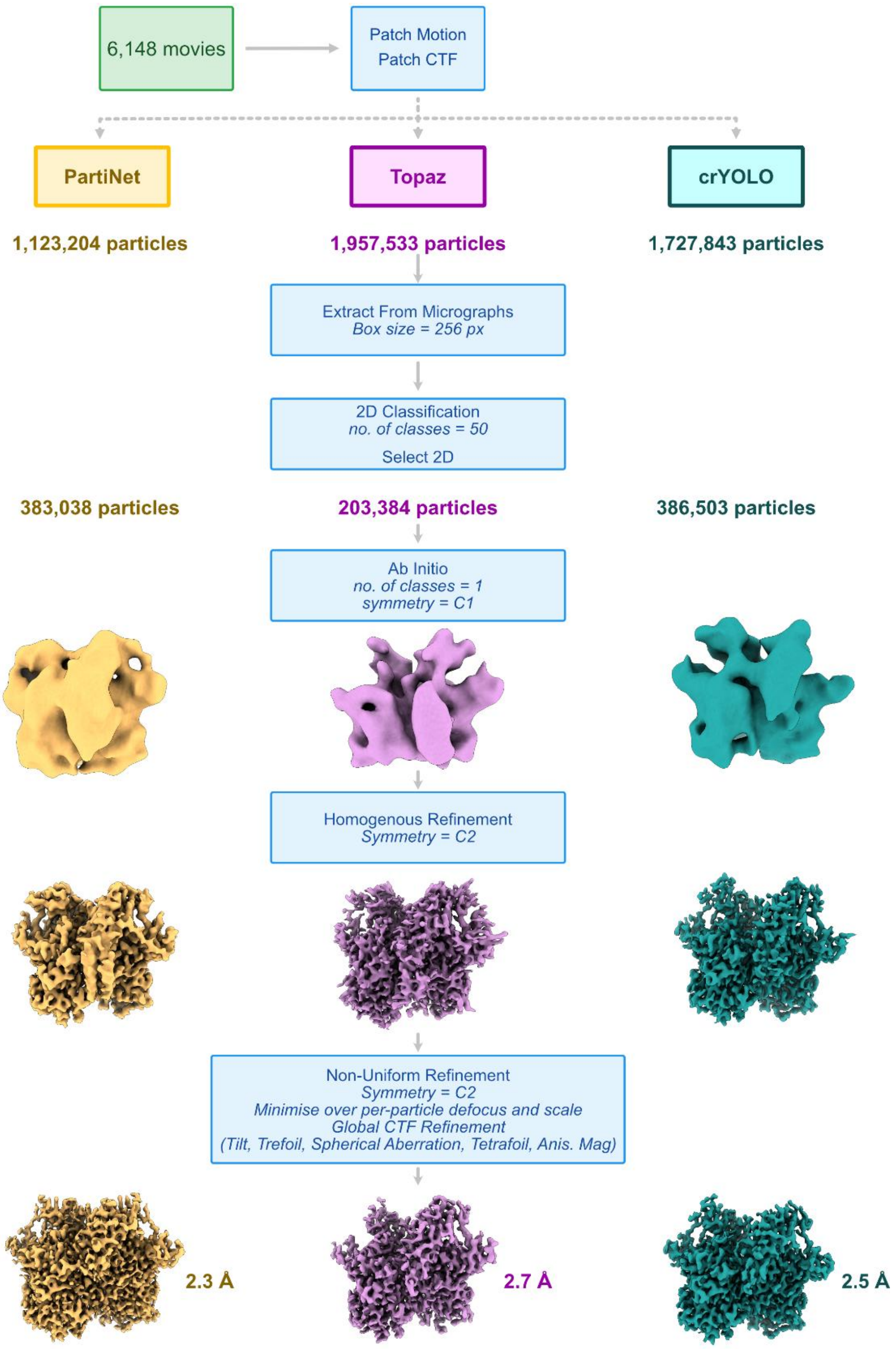


**Supplementary Figure 12.**

**Workflow for processing MORC2**

Details can be found in Methods. Processing was done in CryoSPARC v4.6.2

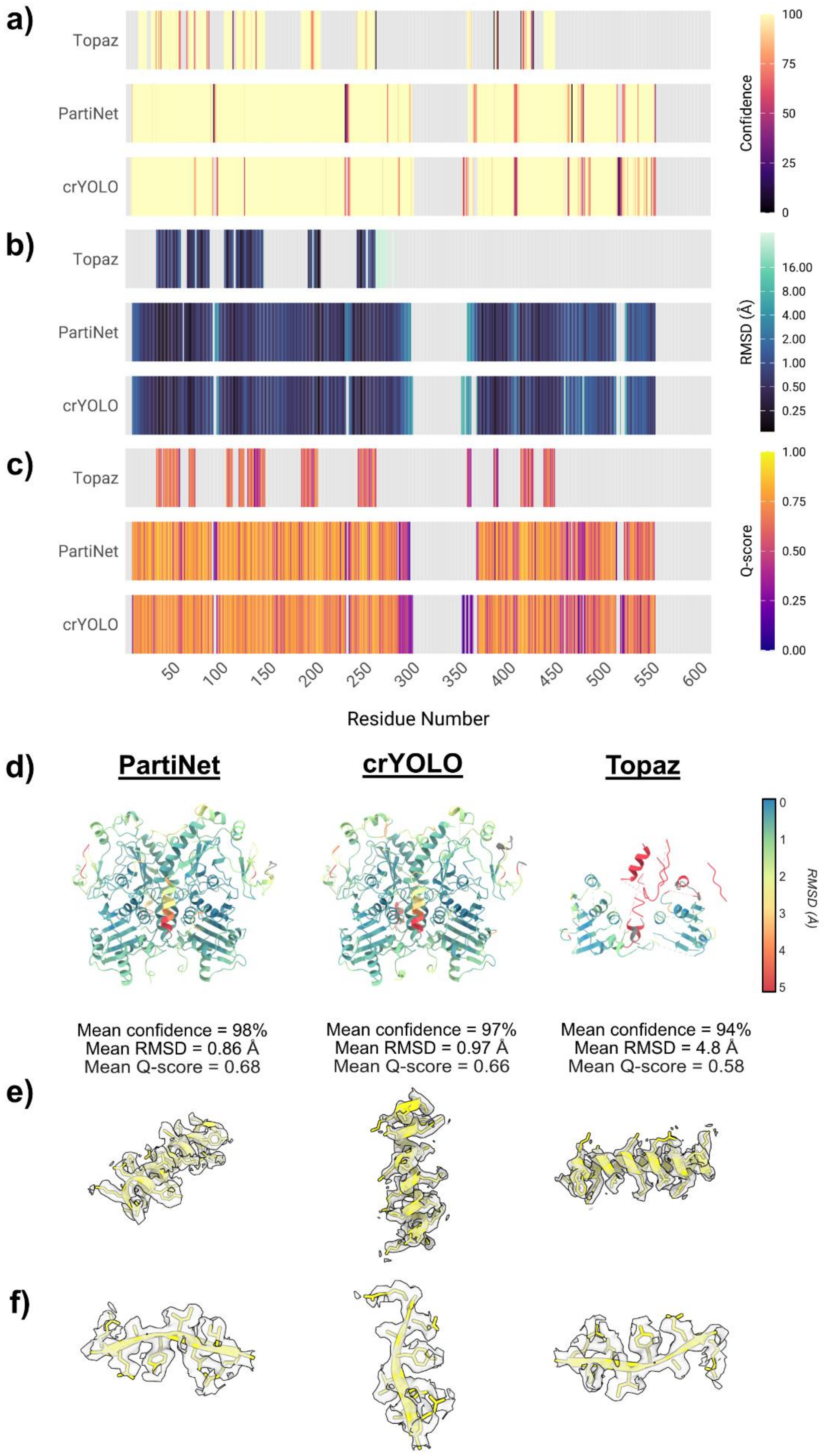
a)
b)
c)
Topaz
PartiNet
crYOLO
Confidence
100
75
50
25
0
RMSD (Å)
16.00
8.00
4.00
2.00
1.00
0.50
0.25
Q-score
1.00
0.75
0.50
0.25
0.00
50
100
150
200
250
300
350
400
450
500
550
600
Residue Number
d)
PartiNet
crYOLO
Topaz
RMSD (Å)
0
1
2
3
4
5
Mean confidence = 98%
Mean RMSD = 0.86 Å
Mean Q-score = 0.68
Mean confidence = 97%
Mean RMSD = 0.97 Å
Mean Q-score = 0.66
Mean confidence = 94%
Mean RMSD = 4.8 Å
Mean Q-score = 0.58
e)
f)

### Supplementary Figure 13.

**ModelAngelo results for MORC2**

**a.** Per-residue confidence of ModelAngelo predictions for MORC2 maps reconstructed with Topaz, PartiNet and crYOLO. **b.** Per-residue RMSD between ModelAngelo prediction and crystal structure of rabbit muscle aldolase (PDB 6ALD). **c.** Per-residue Q-score of ModelAngelo prediction and maps. **d.** Atomic coordinates from ModelAngelo coloured to according to per-residue RMSD for PartiNet, crYOLO and Topaz reconstructions (red indicating high RMSD). Mean confidence, RMSD and Q-score is annotated for each model. **e-f.** Map and model shown for key structural elements.

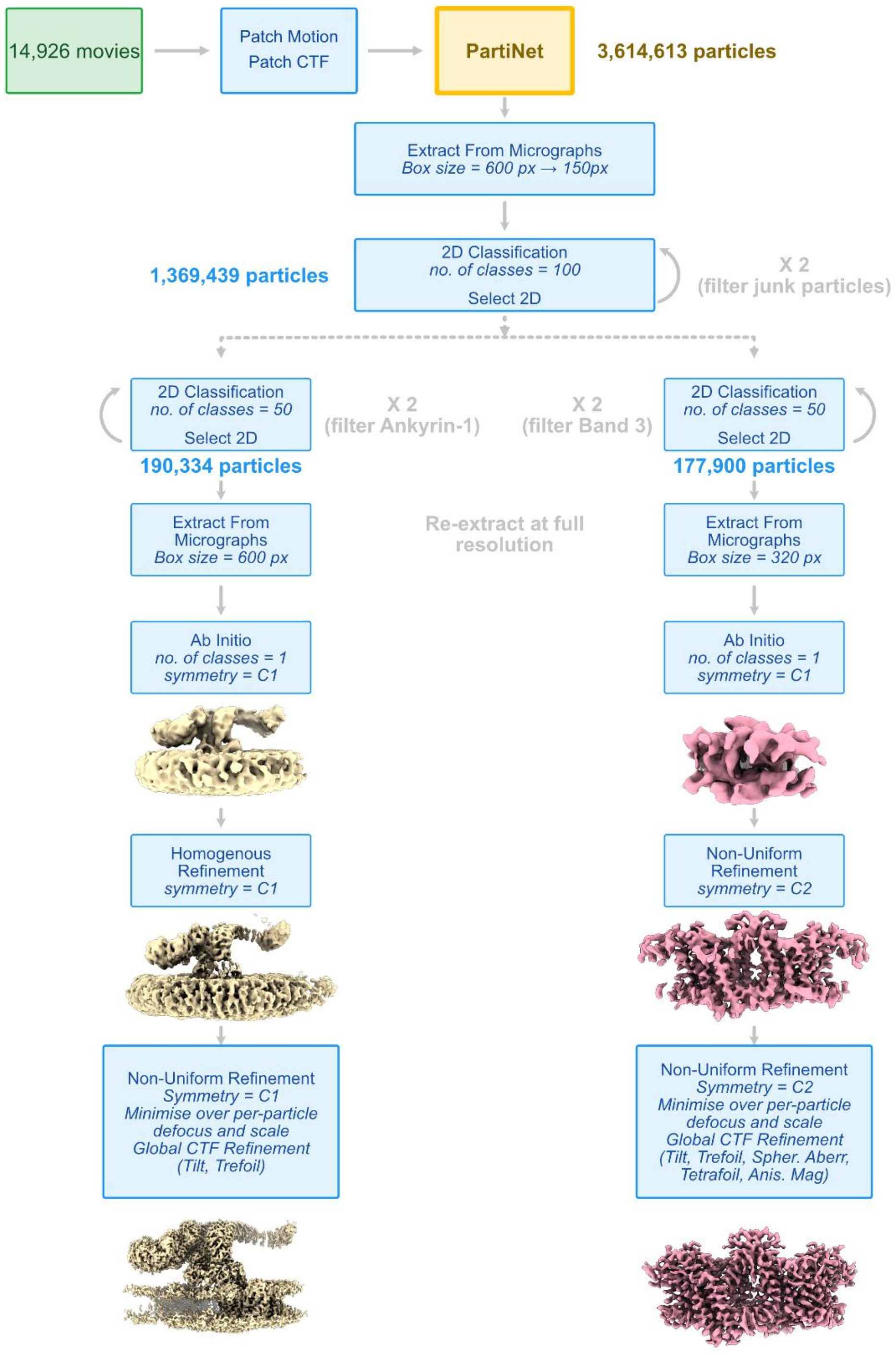


### Supplementary Figure 14.

**Workflow for processing for EMPIAR-11043**

Details can be found in Methods. Processing was done in CryoSPARC v4.7.0

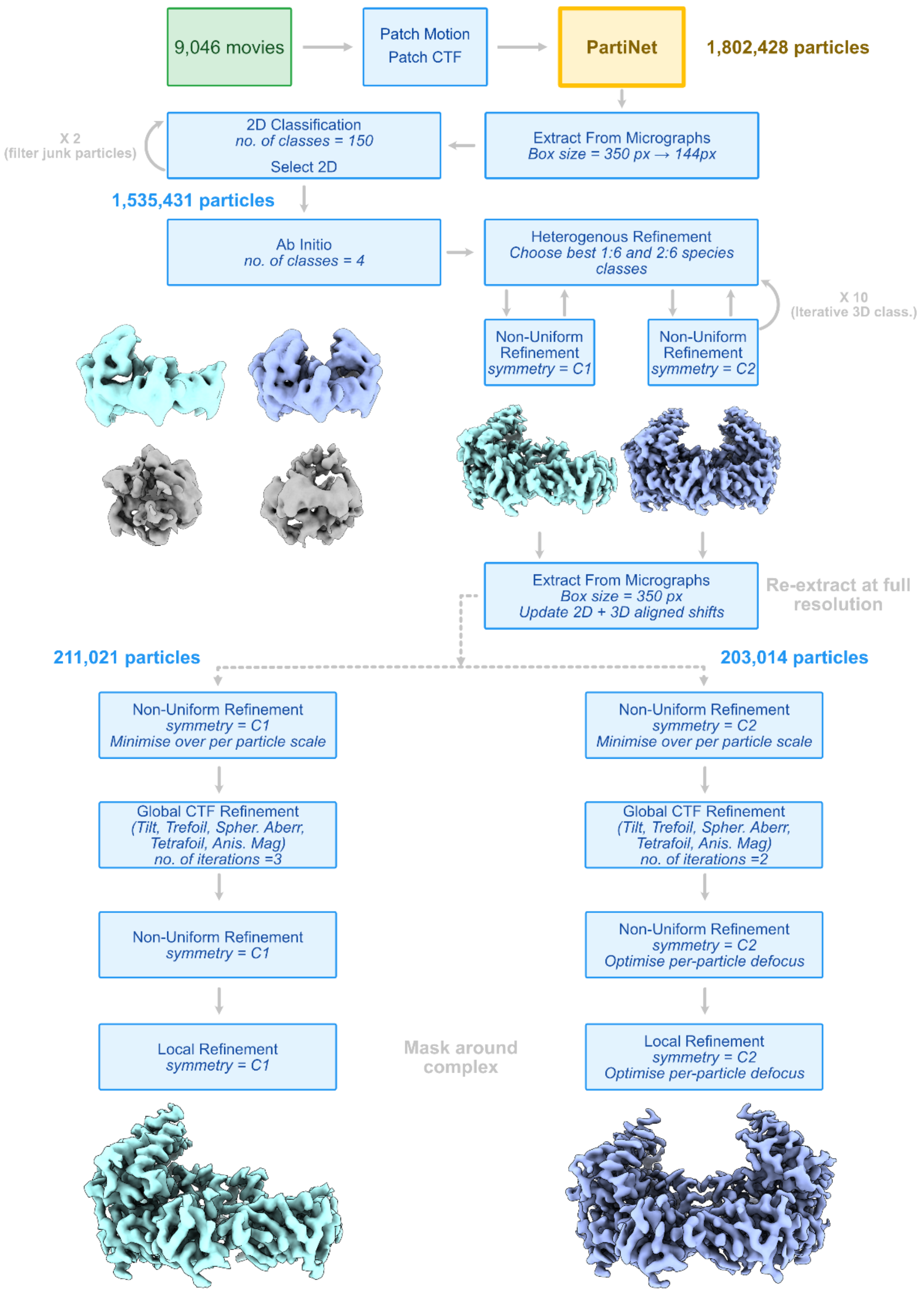


### Supplementary Figure 15.

**Workflow for processing for EMPIAR-12531**

Details can be found in Methods. Processing was done in CryoSPARC v4.7.0

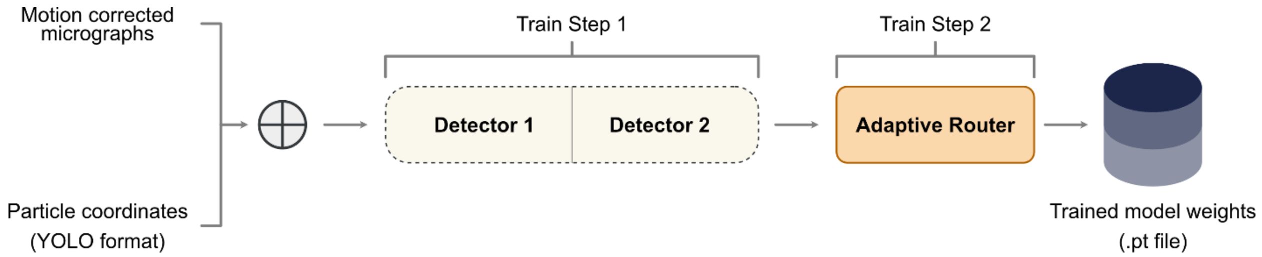


**Supplementary Figure 16.**

**Outline for training PartiNet detailing the two-step process.**

### Supplementary Table 1.

The 34 unique protein datasets were curated by CryoPPP and split into training, validation and test sets

| EMPAIR ID | Protein Type | Size (TB) | Number of Micrographs | Image size | Particle Diameter (px) | Total Structure Weight (kDa) | Number of True Particles |
|---|---|---|---|---|---|---|---|
| 10005 | TRPV1 Transport protein | 0.044 | 30 | (3710, 3710) | 142 | 272.97 | 5374 |
| 10017 | β-galactosidase | 0.005 | 84 | (4096, 4096) | 108 | 450 | 49391 |
| 10028 | Ribosome (80S) | 1.1 | 300 | (4096, 4096) | 224 | 2135.89 | 26391 |
| 10059 | Transport Protein (TRPV1) | 0.062 | 295 | (3838, 3710) | 132 | 317.88 | 190398 |
| 10061 | Hydrolase (Beta-galactosidase) | 0.319 | 300 | (7676, 7420) | 471 | 467.06 | 35218 |
| 10075 | Bacteriophage MS2 | 0.046 | 300 | (4096, 4096) | 233 | 1000 | 12682 |
| 10077 | Ribosome (70S) | 0.774 | 300 | (4096, 4096) | 216 | 2198.78 | 31919 |
| 10081 | Transport Protein | 0.052 | 300 | (3710, 3838) | 154 | 298.57 | 39352 |
| 10093 | Membrane Protein | 0.097 | 300 | (3838, 3710) | 172 | 779.4 | 56394 |
| 10096 | Viral Protein | 1.199 | 300 | (3838, 3710) | 84 | 150 | 231351 |
| 10184 | Aldolase | 0.084 | 300 | (3838, 3710) | 118 | 150 | 219849 |
| 10240 | Lipid Transport Protein | 0.111 | 300 | (3838, 3710) | 156 | 171.72 | 85958 |
| 10289 | Transport Protein | 0.048 | 300 | (3710, 3838) | 162 | 361.39 | 61517 |
| 10291 | Transport Protein | 0.016 | 300 | (3710, 3838) | 130 | 361.39 | 99808 |
| 10345 | Signaling Protein | 0.085 | 300 | (3838, 3710) | 149 | 244.68 | 15894 |
| 10387 | Viral Protein (DNA) | 0.105 | 300 | (3710, 3838) | 213 | 185.87 | 101778 |
| 10389 | Metal Binding Protein | 0.224 | 300 | (3838, 3710) | 313 | 1042.17 | 10870 |
| 10406 | Ribosome (70S) | 0.141 | 300 | (3838, 3710) | 212 | 632.89 | 24703 |
| 10444 | Membrane Protein | 2.399 | 300 | (5760, 4092) | 217 | 295.89 | 58731 |
| 10526 | Ribosome (50S) | 0.46 | 294 | (7676, 7420) | 482 | 1085.81 | 3265 |
| 10532 | Viral Protein | 0.196 | 300 | (4096, 4096) | 174 | 191.76 | 87933 |
| 10576 | Nuclear Protein (DNA) | 0.722 | 295 | (7420, 7676) | 265 | 290.21 | 75220 |
| 10590 | TRPV1 with DkTx and RTX | 0.252 | 300 | (3710, 3838) | 158 | 1000 | 62493 |
| 10669 | Proteasome (Plant Protein) | 13.899 | 300 | (7676, 7420) | 730 | 1681.81 | 19660 |
| 10671 | Signaling Protein | 1.6 | 298 | (5760, 4092) | 133 | 77.14 | 69012 |
| 10737 | Membrane Protein (E-coli) | 0.831 | 293 | (5760, 4092) | 179 | 155.83 | 59265 |
| 10760 | Membrane Protein | 0.199 | 300 | (3838, 3710) | 106 | 321.69 | 173664 |
| 10816 | Transport Protein | 1.5 | 300 | (7676, 7420) | 359 | 166.62 | 45363 |
| 10852 | Signaling Protein | 0.227 | 343 | (5760, 4092) | 123 | 157.81 | 310291 |
| 10947 | Viral Protein | 0.048 | 400 | (4096, 4096) | 240 | 443.92 | 106393 |
| 11051 | Transcription/DNA/RNA | 2.3 | 300 | (3838, 3710) | 214 | 357.31 | 83227 |
| 11056 | Transport Protein | 0.164 | 361 | (5760, 4092) | 164 | 88.94 | 125908 |
| 11057 | Hydrolase | 2.1 | 300 | (5760, 4092) | 186 | 149.43 | 45219 |
| 11183 | Signaling Protein | 0.326 | 300 | (5760, 4092) | 159 | 139.36 | 80014 |

Supplementary Table 2.

**Summary of possible micrograph augmentations during training.**

Values indicate probability of transform being applied to micrograph during training.

| Parameter | Value | Description |
|---|---|---|
| **hsv_h** | 0.015 | HSV hue shift augmentation - randomly adjusts color hue |
| **hsv_s** | 0.7 | HSV saturation augmentation - modifies color intensity/vividness |
| **hsv_v** | 0.4 | HSV value augmentation - adjusts image brightness |
| **translate** | 0.2 | Random translation/shifting of image position |
| **scale** | 0.9 | Random scaling augmentation - resizes images up/down |
| **fliplr** | 0.5 | Horizontal flip probability - mirror image left-to-right |
| **mosaic** | 1.0 | Mosaic augmentation - combines 4 images into one training sample |
| **mixup** | 0.15 | MixUp augmentation - blends two images and their labels |
| **paste_in** | 0.15 | Paste-in augmentation - inserts objects into background scenes |

#### Supplementary Table 3.

**YOLOv7 is published in six different configurations** [2]. As the number of model parameters increases, the reported performance increases and speed of inference (FPS) decreases. A motivation for comparing these configurations was identifying which gave the best performance specifically on cryo-EM micrographs.

| Config | #Param. | Input Size | Reported FPS | Reported mAP@50% |
|---|---|---|---|---|
| YOLOv7 | 36.9M | 640 | 161 | 69.7% |
| YOLOv7-X | 71.3M | 640 | 114 | 71.2% |
| YOLOv7-W6 | 70.4M | 1280 | 84 | 72.6% |
| YOLOv7-E6 | 97.2M | 1280 | 56 | 73.5% |
| YOLOv7-D6 | 154.7M | 1280 | 44 | 74% |
| YOLOv7-E6E | 151.7M | 1280 | 36 | 74.4% |

### Supplementary Table 4.

| | | | | | No. of particles (before Select 2D) | | | No. of particles (after Select 2D) | | | Resolution | | | cFAR | | |
|---|---|---|---|---|---|---|---|---|---|---|---|---|---|---|---|---|
| **EMPIAR ID** | **Protein Name** | **kDA** | **No. of micrographs** | **Defocus range** | **crYOLO** | **Topaz** | **Partinet** | **crYOLO** | **Topaz** | **Partinet** | **crYOLO** | **Topaz** | **Partinet** | **crYOLO** | **Topaz** | **Partinet** |
| **10017** | **Beta-galactosidase** | 450 | 84 | -1.4 to 5.0 | 53,251 | 25,531 | 50,835 | 39,300 | 15,928 | 38,091 | 6.1 | 7.2 | 6.0 | 0.30 | 0.04 | 0.13 |
| **10081** | **HCN1 ion channel** | 299 | 300 | -1.5 to -3.3 | 46,796 | 56,137 | 62,324 | 35,560 | 24,952 | 40,078 | 7.1 | 9.4 | 6.6 | 0.90 | 0.15 | 0.13 |
| **10093** | **NOMPC ion channel** | 779 | 295 | -1.4 to -3.3 | 31,723 | 16,542 | 111,382 | 17,832 | 7,030 | 47,432 | 8.4 | 9.0 | 7.3 | 0.07 | 0.09 | 0.12 |
| **10345** | **Integrin alpha-v beta-8 complex** | 245 | 295 | - | 17,006 | 27,495 | 66,611 | 6,570 | 5,672 | 15,587 | 13 | 15 | 10 | 0.20 | 0.13 | 0.20 |
| **10389** | **Urease** | 1042 | 300 | -0.2 to -0.5 | 11,263 | 18,744 | 140,569 | 7,474 | 6,010 | 11,655 | 7.7 | 7.4 | 6.0 | 0.10 | 0.03 | 0.03 |
| **10532** | **Influenza Hemagglutinin** | 192 | 300 | - | 37,401 | 41,263 | 136,701 | 17,699 | 16,729 | 39,664 | 4.7 | 4.7 | 4.5 | 0.05 | 0.02 | 0.03 |
| **11056** | **NTCP bile acid transporter** | 89 | 305 | 0.8 to 1.6 | 84,614 | 88,315 | 131,851 | 19,422 | 17,529 | 47,692 | 11 | 17 | 8.1 | 0.36 | 0.31 | 0.27 |

### Supplementary Table 5.

**CryoEM data collection, refinement and validation statistics**

| **Data Collection** | **MORC2$^{H3K9\text{-}bound}$ (EMD-68600 and EMPIAR-13226)** |
|---|---|
| Micrographs | 6,140 |
| Particles (Final map) | 383,038 |
| Pixel size (Å) | 0.82 |
| Defocus range (µm) | 0.5 – 2.5 |
| Voltage (kV) | 300 |
| Electron dose (e/Å$^2$) | 60 |
| Symmetry imposed | C2 |
| Initial particle images (no.) | 1,123,204 |
| Final particle images (no.) | 383,038 |
| Resolution (0.143 FSC) (Å) | 2.28 |
| Final Refinement | Non-Uniform (NU) in cryosparc |
| Map sharpening *B* | -98.20 |
| Model used | de novo using ModelAngelo |
| Unmodelled regions (in comparison to crystal structure) | 308-342 |
| $CC_{map\ model}$ | 0.78 |
| **Model quality** | |
| Bond length (Å) / Bond angles (°) | 0.003/0.556 |
| **Ramachandran** | |
| Favoured (%) | 96.28 |
| Outliers (%) | 0.11 |
| Rotamer outliers (%) | 1.45 |
| C-Beta deviations (%) | 0.00 |
| Clashscore | 7.19 |
| MolProbity Score | 1.74 |
